# An Integrative Approach to Develop and Characterise Antibodies Against the Cancer Associated Antigen Sialyl Lewis A (CA 19-9)

**DOI:** 10.1101/2025.06.15.659274

**Authors:** Anika Freitag, Sana Khan Khilji, Ruslan Nedielkov, Shalini Murali Kumar, Michael Krummhaar, Janine Arndt, Gustavo Marçal Schmidt Garcia Moreira, Jost Luhle, Felix Goerdeler, Carsten Kamphues, Maria Andrea Mroginski, Christian Roth, Peter H. Seeberger, Heiko M. Moller, Oren Moscovitz

## Abstract

**Background:** Sialyl Lewis A (sLeA), or the CA 19-9 marker, is a tetrasaccharide and a tumour-associated carbohydrate antigen (TACA) overexpressed and abnormally secreted as a serum-borne marker in gastrointestinal malignancies. CA 19-9 is the best validated and only FDA-approved serologic marker clinically used to monitor recurrence, progression, and therapy efficiency in pancreatic ductal adenocarcinoma (PDAC) patients. Due to its altered expression on cancer cells, sLeA is also an attractive target for antibody development. Although recent clinical trials have demonstrated insufficient efficacy of the fully human anti-sLeA 5B1 (MVT-5873) format as a stand-alone drug or an adjuvant therapy in PDAC [1], its safety profile and unique expression in additional malignancies keep CA 19–9 an attractive TACA. Hence, we set out to explore the use of synthetic sLeA to develop novel monoclonal antibodies (mAbs) with improved sLeA recognition and better efficacy.

**Methods:** Two mAbs targeting sLeA were generated through mice immunisation with synthetic sLeA glycoconjugates, synthetic glycan arrays, and hybridoma technology. We then compared the antigen-binding properties of the newly developed mAbs with the widely used mAb 1116-NS-19- 9 via synthetic glycan arrays, immunohistochemistry (IHC), X-ray crystallography, molecular dynamics (MD) simulation, and Saturation Transfer Difference Nuclear Magnetic Resonance (STD NMR) spectroscopy.

**Results:** The newly generated mAbs demonstrated improved affinity and specificity for both synthetic and native sLeA, surpassing the performance of the established mAb 1116-NS-19-9. First, synthetic glycan arrays, surface plasmon resonance (SPR), and isothermal titration calorimetry (ITC) assays confirmed superior antigen-binding properties to synthetic sLeA. In particular, the mAb designated GB11 demonstrated markedly enhanced binding to native sLeA ectopically expressed in B16 melanoma cells. To elucidate the structural origin of GB11’s improved antigen binding, we conducted high-resolution mapping of the molecular recognition patterns between sLeA and the different antibodies using X-ray crystallography and STD NMR. These analyses revealed subtle yet critical differences in the glycan engagement and identified key structural features underlying GB11’s enhanced recognition of sLeA. MD simulations further supported these observations, indicating distinct orientations of sLeA within the binding pockets of each mAb.

**Conclusion:** Our results suggest better recognition of the sLeA antigen by the newly generated GB11 antibody and provide a detailed high-resolution elucidation of the molecular interactions behind it. Our study may provide a novel tool with improved theranostic properties against sLeA-overexpressing malignancies.

## Background

Glycosylation is the most abundant form of protein post-translational modification taking part in every cellular process as part of the glycocalyx, including cell-cell adhesion, cell-matrix interaction, immune recognition, and membrane organisation, among many others. It is not template-driven but mediated by coordinated functions of glycosyltransferases and glycosidases and takes place in the endoplasmic reticulum and Golgi compartment of eukaryotic cells. Glycosylation varies across cells and tissues and depends on the cell microenvironment, causing structural complexity and heterogeneity in glycans [2–4]. Unsurprisingly, altered glycosylations are a hallmark of many diseases, including cancer. These aberrant tumour-associated carbohydrate antigens (TACAs) can, therefore, be used to distinguish cancer from healthy cells. TACAs are not only representative of the changes in neoplastic cell behaviour but also play essential roles in the development, metastatic progression, and immune system evasion of cancer [5–7]. Due to their abnormal structure or expression level on cancer cells, TACAs can be very attractive candidates for vaccine and antibody-targeted therapy [8]. However, owing to their enormous diversity and structural complexity, obtaining enough homogeneous material of naturally derived glycans to immunise animals with and produce specific antibodies (Abs) against poses a considerable challenge [9–11]. While total chemical synthesis can provide pure, well-defined glycans, it is laborious and time-consuming. The development of automated glycan assembly has accelerated and simplified the process significantly [12].

Sialyl Lewis A (sLeA), also known as the CA 19-9 marker, is a tetrasaccharide composed of Neu5Ac(α2-3)Gal(β1-3)[Fucα(1-4)]GlcNAc. Sialyl LeA is part of the human histo-blood system, known as the Lewis antigen system, which consists of the type I and type II Lewis antigens [13]. H1, H2, Lewis A (LeA), LeB, LeX, and LeY share the same three monosaccharides, namely N-Acetylglucosamine (GlcNAc), Galactose (Gal), and Fucose (Fuc), differing only in their corresponding glycosidic bonds (type I: Gal(β1-3)GlcNAc; type II: Gal(β1-4)GlcNAc). Further addition of sialic acid (N-Acetylneuraminic acid (Neu5Ac)) to LeA and LeX leads to sLeA and sLeX, respectively [13].

While it is found in embryonic tissue and is present at low levels in healthy tissue, sLeA synthesis is upregulated in specific epithelial cancers, including digestive tract, liver, breast, lung, pancreatic, and ovarian cancer, as well as non-cancer diseases, such as diabetes mellitus, obstructive jaundice, and pancreatitis (which can predispose to pancreatic cancer) [14–17]. Moreover, it stands as the only US Food and Drug Administration (FDA)-approved biomarker for monitoring pancreatic cancer progression, which is generally known for its exceptionally low 5- year survival rate (9%) and poor prognosis (∼5 months) post-diagnosis [18–21].

Sialyl LeA interacts with E- or P-selectins present on endothelial cells, thereby facilitating tumour vascularisation during metastasis and allowing cancer cells to migrate to the metastatic site [15]. In healthy tissue, disialyl Lewis A interacts with the immunosuppressive siglec-7 and -9 receptors on macrophages/monocytes and CD8^+^ T cells to help maintain immunological homeostasis. However, elevated expression of sLeA abrogates this interaction, leading to chronic inflammatory stimuli that may support cancer progression [22]. Furthermore, abnormal sLeA expression promotes inflammation by recruiting circulating lymphocytes to peripheral lymph nodes and cancerous tissue [23]. Reciprocally, an inflammatory environment can enhance the expression of tumour-associated sialylated antigens such as sLeX/A by pro-inflammatory cytokines that regulate the expression of sialyltransferases [24].

The Database of Anti-Glycan Reagents (DAGR) currently lists 43 Abs targeting sLeA [25], including the monoclonal antibodies (mAbs), murine 1116-NS-19-9 and human 5B1, that are most commonly used for clinical diagnosis and therapy of pancreatic cancer, respectively [26]. The cytotoxic 5B1 (now MVT-5873 or BNT321) is a high-affinity human mAb that was generated from blood lymphocytes of a breast cancer patient immunised with synthetic sLeA conjugated to KLH [27], and is currently part of several ongoing or recently finished clinical trials [28–32]. 1116-NS- 19-9 is an IgG1 that was generated in mice in the early 1980s by immunising with five different human colorectal carcinoma (CRC) cell lines [32–34]. How carbohydrate homogeneity and mode of presentation during the immunisation process affects the finally recognised carbohydrate epitope and the affinity of carbohydrate-specific antibodies is still not fully understood. To evaluate how the “natural” immunisation caused by aberrantly glycosylated cancer cells (such as in the case of 1116-NS-19-9) compares to the immunisation with a well-defined carbohydrate antigen, we used a construct of synthetic sLeA conjugated to CRM197 (sLeA- CRM_197_) to immunise mice and isolated two murine antibodies, namely GB11 and HA8, which we characterised and compared to the murine 1116-NS-19-9.

## Methods

### Conjugation of sialyl Lewis A and CRM_197_

3’-sialyl Lewis A- (sLeA) was purchased from GlycoUniverse GmbH. CRM_197_ was conjugated to sLeA following standard protocols [35,36]. Briefly, 1 eq. of sLeA was mixed with 10 eq. of homobifunctional bis(*p*-nitrophenyl) adipate in 300 µL DMSO + 25 µL pyridine + 10 µL triethylamine (TEA) and stirred for 3 h at 300 rpm, room temperature (RT). After lyophilisation, the glycan half ester was washed 3x with chloroform and 3x with dichloromethane until uncoupled linker was no longer detected in the wash fractions. Next, 70 eq. of washed glycan half ester were mixed with 1 eq. of CRM_197_ in 140 µL conjugation buffer (0.1 M sodium phosphate pH 8) and stirred for 24 h at 70 rpm, RT. The degree of loading (DOL) was assessed by MALDI in linear positive ion mode using a 2,5-DHAP matrix and an Autoflex Speed (Brucker Daltonics). Comparing spectra of unconjugated CRM_197_ and sLeA-CRM_197_, we determined a DOL of 7.4 glycans/protein (Figure S1).

### Mouse Immunisation

Equal volumes of sLeA-CRM_197_ (in sterile PBS) and aluminum hydroxide Al(OH)_3_ (Alhydrogel) were mixed overnight at 4°C to allow adsorption of the glycoconjugate to the adjuvant matrix. Five 6- to 8-week-old female C57BL6JRj mice were immunised subcutaneously with 1 µg sLeA (equal to 8.65 µg sLeA-CRM_197_) per injection. The mice received the glycoconjugates 4 times on day 0, 14, 28, and 42, followed by a boost injection on day 68. Three days after the boost injection, the spleen were harvested. Animal experiments were performed by Hybrotec GmbH (Germany, Potsdam) and approved by the Landesamt fur Arbeitsschutz, Verbraucherschutz und Gesundheit (LAVG) Brandenburg (Gesch-Z. 2347-A-34-1-2020). Experiments were performed according to the German law, following the regulations of the Society for Laboratory Animal Science (ALAS) and of the Federation of Laboratory Animal Science Associations (FELASA).

### Monoclonal Antibody Development

Monoclonal antibodies were obtained via hybridoma technology from mouse splenocytes as previously described in Broecker et al. [35]. The selection of hybridoma clones was done by glycan microarray-assisted analysis [10]. After three subsequent subcloning steps, two hybridoma clones, GB11 and HA8, producing Immunoglobulin G1 (IgG1) exclusively binding to sLeA were recovered.

### Hybridoma Culture

Hybridoma cells were stored at -80°C in RPMI + 10% FCS and 10% DMSO at a concentration of 2x10^6^/mL. For a new culture, cells were thawed at 37°C for 30 s before they were washed in ISF- 1 medium with 10% FCS to remove DMSO. Cells were centrifuged at 300 g/RT for 5 min. The supernatant was removed and taken up afresh in 10 mL of ISF-1 + 10% FCS. The cells were cultured at 37°C in the presence of 5% CO_2_ and the FCS was gradually decreased over days until it was completely removed. Cells were then passaged every 2-3 days until they were expanded to 300 cm^2^ flasks. They were then allowed to grow for 7-10 days until they died. The cultures were centrifuged at 1,200 g/RT for 20 min. The supernatant containing mAb was sterile filtered with a 0.2 µm filter and stored with 0.02% NaN_3_ at 4°C to avoid contamination.

### RNA Isolation and Sequencing of the mAb GB11 and HA8

RNA was isolated from 10^7^ hybridoma cells of either clone GB11 or HA8, using RNeasy MiniKit (Qiagen, Hilden, Germany). The cDNA of the variable domains was amplified via template-switch reverse transcription as described elsewhere [37]. Briefly, cDNA was synthesised using chain-specific primers for mouse IgG heavy chain, kappa light chain, and lambda light chain in the presence of a template-switch oligo (AAGCAGTGGTATCAACGCAGAGTACATrGrGrG, where r stands for RNA base) to add a 3’ adaptor sequence for subsequent sequencing. Reverse transcription was performed using the SuperScript IV Reverse Transcriptase system (Invitrogen), which is capable of template switching. Subsequently, cDNA from each chain was amplified via PCR using chain-specific nested reverse primers that were modified with 5’-phosphates, along with an adaptor sequence-specific universal forward primer. Generated fragments were purified on a 2% agarose gel and blunt end ligated into the pCRZeroT plasmid, previously digested with SmaI (NEB). The pCRZeroT plasmid was a gift from Ken Motohashi (Addgene plasmid #120276; RRID: Addgene_120276 [38]). Ligation products were transformed into the *Escherichia coli* (*E. coli*) DH5alpha strain. Plasmid DNA from successfully transformed colonies was isolated and sequenced using the M13 standard primers to obtain variable domain sequences.

### Monoclonal Antibody Purification

Fast protein liquid chromatography (FPLC) was performed at 4°C on an AKTApurifier UPC10 System (GE Healthcare; Uppsala, Sweden) operated with the UNICORN 5.11 software. A 5 mL Pierce Protein A/G Chromatography cartridge (Thermo Scientific; Rockford, IL, USA) was equilibrated with binding buffer (25 mM Sodium phosphate 150 mM NaCl, pH 7.4) by passing through 10x column volumes (CV) through the cartridge. Up to 2 L of hybridoma supernatant containing the mAb was passed through the column overnight, at flow rates ranging from 0.5 to 1 mL/min. The cartridge was washed with 15xCV binding buffer followed by 10xCV of a mixture of 80% binding buffer and 20% protein A/protein G elution buffer (100 mM Glycine-HCl, pH 2.7). Finally, the resin-bound mAb was eluted with 15-20xCV 100% protein A/protein G elution buffer at a flow rate of up to 5 mL/min and collected in 5 mL fractions, containing a previously measured amount (approx. 155 µL) of protein A/protein G neutralisation buffer (1 M Tris-HCl, pH 9), for a final pH of 7.4. The presence of protein within the fractions was followed by UV absorption at 280 nm. The fractions containing eluted mAb were concentrated using a centrifugal Amicon® Ultra-4 filter (MWCO 30 kDa) (Merck Millipore; Tullagreen, Ireland) and further purified by size exclusion chromatography on the AKTApurifier with a HiLoad 16&600 Superdex 200 pg (S200) (Cytiva) in 1x phosphate-buffered saline (PBS, pH 7.4). All fractions were analysed by SDS-PAGE. The eluted fractions containing the mAb were concentrated using a centrifugal Amicon® Ultra-4 filter (MWCO 30 kDa), and the protein concentration was determined with a NanoDrop ND-1000 Spectrophotometer (Thermo Scientific; Waltham, MA, USA). The mAb was stored in 1xPBS containing 0.02% NaN_3_ at 4°C.

### Thermal Shift Assay

The mAbs GB11, HA8, and 1116-NS-19-9 were diluted with PBS to 0.15 mg/mL. Intrinsic fluorescence was recorded at 330 nm and 350 nm while heating the sample from 35 to 95°C at a rate of 3°C/min. For the data collection as well as calculation of the ratio of fluorescence (350/330 nm) and the inflection temperature T*i*, the NanoTemper Tycho NT.6 (NanoTemper Technology GmbH, Munich, Germany) was used according to the manufacturer’s instructions.

### Isothermal Titration Calorimetry

ITC analyses were conducted on an iTC200 instrument (MicroCal). GB11, HA8, and 3’- sialyl Lewis A (BIOSYNTH Carbosynth, OS00745) were diluted in PBS buffer, pH 7.4. 300 µL of 12.5-20 µM mAb was loaded into the sample cell, while sLeA was titrated with the syringe via 18 injections. The first injection was of 0.4 µL, while the rest were of 2 µL each. The final molar ratio of mAb to sLeA was in both cases between 1:20 and 1:24. All measurements were conducted at 25°C. Data was analysed with MicroCal PEAQ-ITC Analysis Software. For both mAbs, at least three repetitions were carried out. Due to the financial constraints associated with purchasing the 1116-NS-19-9 in amounts needed to carry out triplicates, this mAb was not included as a positive control in our analysis. Instead, the glycan was titrated to 1x PBS buffer as a negative control.

### Surface Plasmon Resonance

Binding experiments were carried out on a Biacore T100 instrument (Cytiva Life Sciences, Danaher Corporation) using the Biacore T200 control software. On a CM5 chip, the Mouse Antibody Capture Kit (Cytiva Life Sciences, Danaher Corporation) was used according to manufacturer’s protocol to first immobilise ∼3,000-5,000 RU of α-mouse IgG capture antibody on two flow cells, one of which was used for mAb immobilisation and the other as a ‘blank’- immobilised flow cell for reference to compensate for unspecific binding of glycan to sensor chip surface and α-mouse IgG capture antibody. Immobilisation of mAbs and glycan binding assays were performed at 25°C in PBS. Approximately 400 RU of mAbs were captured at either a concentration of 50 µg/mL for IgG GB11 and HA8 or a 1:100 dilution factor for 1116-NS-19-9 (Thermo Fisher #MA5-12421). The surface contact time was 180 s with a flow rate of 30 µL/min and a stabilisation period of 60 s to allow any uncaptured mAb to be washed away. Following this, sLeA was run over the immobilised mAbs in 2 cycles with increasing concentrations ranging from 0.375 µM to 100 µM so that 10 different concentrations could be measured. The highest concentration of the first cycle was taken as the lowest concentration of the second, and both cycles were plotted on the same graph using “Single cycle kinetics”. The parameters used were a contact time of 60 s and a dissociation time of 180 s at a flow rate of 30 µL/min. Flow cells were regenerated with 10 mM glycine-HCl pH 1.7 with a contact time of 120 s at a flow rate of 20 µL/min, and a stabilisation time of 300 s was added to ensure that the baseline returned to its original value and all secondary mAb molecules were removed. For all mAbs, 3 repetitions were carried out. Analyses were carried out using the Biacore T200 evaluation software 3.2. As the on- and off-rates were outside of the measurable ranges of the instrument, the K_D_ values were determined using a “steady-state affinity” model. Each replicate dataset for every mAb was normalised such that the highest recorded response unit (RU) was set to 100%. The normalised binding responses were then plotted against the logarithm of sLeA concentrations. Curve fitting was performed using the “log(agonist) vs. response – variable slope” model in GraphPad Prism (v10.4.2, Windows), applying minimum and maximum constraints of 0 and 100. The resulting data were fitted to the following equation:

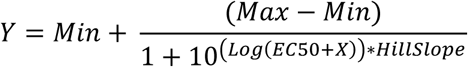

X: log of concentration; Y: Response, increasing as X increases; Minimum (Min) and Maximum (Max): Plateaus in same units as Y (Min = 0; Max = 100); logEC50: same log units as X; Hill Slope: Slope factor or Hill slope, unitless.

### Glycan Array Assay

The synthetic glycans (0.1 mM) were printed in lab on glass slides as described elsewhere [39,40]. The arrays contained immobilised sLeA and structurally related glycans of the Lewis family such as LeA and B, LeX and Y, sLeX, as well as additional TACAs like sialyl Tn (sTn) and Tn. As negative controls, the glycan array included immobilised protein (100 µg/mL) such as a mouse IgG and CRM_197_ (Table 1, Figure S7). The printed slide was blocked with 50 µL of 3% BSA/PBS per well at 37°C. After 1 h incubation, the wells were washed once with 50 µL PBS and incubated for 1 h at 37°C/slight shaking with 50 µL of 5 µg/mL of either HA8, GB11 or 1116-NS- 19-9 (OriGene Technologies, Inc., #CF190083) in 1% BSA/PBS. After the incubation of the primary mAb, the wells were washed twice with 50 µL PBS for 15 min at 37°C/slight shaking each. The secondary antibody Alexa Fluor^TM^ 635, goat α-mouse-IgG (H+L) (Invitrogen by Thermo Fisher Scientific) was added with a dilution of 1:500 in 1% BSA in PBS and incubated for 1 h at 37°C. The wells were then washed twice with 50 µL PBS for 5 min each. As a final wash, the 64- well grid was removed, and the slide was dipped several times into ddH2O in a 50 mL falcon. Once dried by centrifugation at 300 g for 5 min, the slide was directly scanned using a Glycan Array Scanner Axon GenePix® 4300A (Molecular Devices, LLC, San Jose, CA, USA). The binding was analysed using GenePix Pro7 (Molecular Devices, LLC).

**Table 1).** Glycans printed on CodeLink® – glass slide. The table lists the synthetic glycan structures that were printed on the slide with a concentration of 0.1 mM in a humidity chamber at RT. The numbering correlates with the numbering of the printing pattern found in Figure S8a as well as their SNFG representation in Figure S8e.

| No | Printed Compound | Carbohydrate Structure |
| --- | --- | --- |
| 1 | Lewis A - (amino linker) | Gal(β1-3)[Fuc(α1-4)]GlcNAc(β1-1)aminopentanol |
| 2 | Lewis B - (amino linker) | Fuc(α1-2)Gal(β1-3)[Fuc(α1-4)]GlcNAc(β1-1)aminopentanol |
| 3 | Lewis X - (amino linker) | Fuc(α1-3)[Gal(β1-4)]GlcNAc(β1-1)aminopentanol |
| 4 | Lewis Y - (amino linker) | Fuc(α1-3)[Fuc(α1-2)Gal(β1-4)]GlcNAc(β1-1)aminopentanol |
| 5 | Sialyl Lewis X - (amino linker) | Fuc(α1-3)[Neu5Ac(α2-3)Gal(β1-4)]GlcNAc(β1-3)-Gal(β1-4)Glc(β1-1)aminohexanol |
| 6 | **Sialyl Lewis A - (amino linker)** | **Neu5Ac(α2-3)Gal(β1-3)[Fuc(α1-4)]GlcNAc(β1-1)aminopentanol** |
| 7 | Lactose - (amino linker) | Gal(β1-4)Glc(β1-1)aminopentanol |
| 8 | Sialyl Tn - (amino linker) | Neu5Ac(α2-6)GalNAc(α1-1)aminopentanol |
| 9 | Tn - (amino linker) | GalNAc(α1-1) |
| 10 | H-antigen type 2 - (amino linker) | Fuc(α1-2)Gal(β1-4)GlcNAc(β1-3)Gal(β1-4)Glc(β1-1)aminopentanol |
| 11 | Linear Hexamer - (amino linker) | Gal(β1-4)GlcN(β1-3)Gal(β1-4)GlcN(β1-3)Gal(β1-4)Glc(β1-1)aminopentanol |
| 12 | GM1b (cisGM1) - (amino linker) | Neu5Ac(α2-3)Gal(β1-3)GalNAc(β1-4)Gal(β1-4)Glc(β1-1)aminopentanol |
| 13 | GM3 - (amino linker) | Neu5Ac(α2-3)Gal(β1-4)Glc(β1-1)aminopentanol |
| 14 | Sialyllacto-N-tetraose - (amino linker) | Neu5Ac(α2-3)Gal(β1-3)GlcNAc(β1-3)Gal(β1-4)Glc(β1-1)aminopentanol |
| 15 | SLPG (S-i, nHM1) - (amino linker) | Neu5Ac(α2-3)Gal(β1-4)GlcNAc(β1-3)Gal(β1-4)GlcNAc(β1-3)Gal(β1-4)Glc(β1-1)aminopentanol |
|  | Mouse IgG | Unrelated mouse IgG to confirm secondary binding |
|  | CRM<sub>197</sub> | Carrier protein during immunisation |

### Immunofluorescence Staining

Surgically resected tissue was fixed in neutral buffered 4% formaldehyde for 16-24 h and processed for paraffin embedding. After deparaffinisation of 4 μm tissue sections, antigen unmasking was performed using 10 mM sodium citrate buffer (pH 6.0) with 0.05% Tween for 60 min at 90°C. Paraffin sections were permeabilised with 0.5% Triton X-100 for 10 min and blocked with 5% normal goat serum and 1% BSA for 60-120 min. Incubation with primary mAbs was performed overnight at 4°C in blocking solution using either the commercial mouse mAb, 1116-NS-19-9 (1:100, Thermo Fisher #MA5-12421), or the newly generated mouse mAb, GB11 or HA8 (5 µg/mL), against sLeA, simultaneously with a rabbit mAb against Vimentin (1:250, clone EPR3776, Abcam #ab9254). After rinsing with PBS, sections were incubated for 60-90 min at 37°C in 1% BSA with 4′,6-diamidino-2-phenylindole (DAPI, 1:1,000) and the secondary mAbs goat anti-mouse Alexa Fluor^TM^ 594 (1:250**, Thermo Fisher** #**A-11005**) as well as goat anti-rabbit Alexa Fluor^TM^ 488 (1:500, Abcam #ab150081). The paraffin sections were embedded in ProTaqs® MountFluor and analysed using the confocal Laser Scanning Microscope 510 META (Zeiss).

### Cell Culture

B16 and B16FUT3+ cells were kindly gifted by Prof. Dr. J. V. Ravetch [41]. The cells were grown in Dulbecco’s Modified Eagle Medium (DMEM; PAN^TM^ Biotech), supplemented with 10% foetal calf serum (FCS, PAN^TM^ Biotech), 2 mM/mL L-glutamine (PAN^TM^ Biotech), 10 U/mL penicillin/10 µg/mL streptomycin (PAN^TM^ Biotech) at 37°C in the presence of 5% CO_2._ For B16 FUT3+, 500 µg/mL of Geneticin (Gibco^TM^) as a selection antibiotic were additionally added to the growth medium. For binding assays, cells were collected out of culture flasks using Trypsin/EDTA.

### Flow Cytometry

1 x 10^6^ cells per sample were used. After centrifugation for 5 min at 300 g, the cell pellet was incubated with 5 µg/mL of either HA8, GB11, or 1116-NS-19-9 (OriGene Technologies, Inc., #CF190083) in PBS for 1 h at 25°C. For the titration binding assays, a concentration range of 0.155 ng/µL – 40 ng/µL for all mAbs was used. The cell pellet was then washed twice with PBS, after a centrifugation of 5 min at 300 g. Following the washing steps, the cells were incubated for 1 h with Alexa Fluor^TM^ 635, goat α-mouse-IgG (H+L) (Invitrogen by Thermo Fisher Scientific) (1:500 dilution in PBS). The samples were then washed three times with PBS, resuspended in 200 µL PBS and analysed by flow cytometry.

The data was collected on a FACSCantoII (BD Biosciences, San Jose, CA, USA) and analysed with FlowJo (v10.8.1, FlowJo LLC, Ashland, OR, USA). First, intact cells were selected (FSC-A over SSC-A) and then gating was carried out for single cells (FSC-W over FSC-H). The binding of mAb was examined in histograms of the APC channel.

The apparent K_D_ values were determined by normalising the measured mean fluorescence intensity (MFI) to the highest MFI observed across five independent experiments for each antibody. The normalised MFI values were then plotted against the logarithmic concentrations of the mAbs. Curve fitting was performed using the “log(agonist) vs. response – variable slope” model in GraphPad Prism (v9.3.1, Windows), with the bottom and top constraints set to 0 and 100, respectively. The EC50 value, defined as the mAb concentration required to elicit half-maximal response, was used to determine apparent binding affinity. The fitted data were modelled using the following equation:

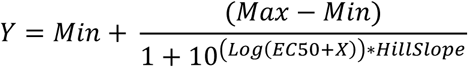

X: log of concentration; Y: Response, increasing as X increases; Minimum (Min) and Maximum (Max): Plateaus in same units as Y (Min = 0; Max = 100); logEC50: same log units as X; Hill Slope: Slope factor or Hill slope, unitless.

### Ficin Digestion

For GB11 and HA8, after purification, the eluted and concentrated mAb was injected onto an S200 column for buffer exchange to 100 mM sodium citrate/6.36 mM EDTA pH 6. Once concentrated, 4.39 mg/mL end concentration cysteine was added and incubated overnight with Ficin immobilised on agarose beads. The Fab product from the digestion was purified from the mixture with a 5 mL Pierce Protein A/G Chromatography cartridge (Thermo Scientific; Rockford, IL, USA) and buffer exchanged to the crystallisation buffer 10 mM Tris-HCl pH 7, 150 mM NaCl with the S200 column.

### Crystallisation, Data Collection, and Structure Solution

Crystals of Fab fragments were grown using vapour diffusion in a hanging drop. Initially, 4x 96- well plates were set up with various crystallisation conditions. Sitting drops of 0.2 µL were added to the wells using an Oryx4 pipetting robot (Douglas Instruments). The plates were stored at RT until crystals were seen. The conditions giving the best crystals were set up in 48 well plates (condition 1: 2.06 M DL-malic acid pH 7.0, 2 mM citric acid pH 3.5, 0.5% PEG3350; condition 2: 1.6 M sodium malonate pH 7, 30 mM Hepes pH 7.0, 8.3% PEG3350). At a concentration of 12.4 mg/mL and 9.8 mg/mL, 2 µL of Fab fragments of GB11 or HA8, respectively, were mixed in a 1:1 ratio in the two conditions mentioned above. Crystals typically appeared within a week, and emerging crystals were fished. HA8 aggregated in the preliminary crystallisation steps and did not give crystals. Apo crystals of GB11 were directly frozen in liquid nitrogen, while holo crystals were incubated for seven days in mother liquor supplemented with 5 mM sLeA and then frozen in liquid nitrogen.

Diffraction data were collected at Berlin BESSY II, beamline 14.1 and 14.2 at 100 K. Data were processed with Xia2/DIALS [42–45], and the structure of GB11 without ligand was solved by molecular replacement with coordinates of 1116-NS-19-9 Fab fragment (PDB ID 6XTG) using MR-Phaser [46]. The structure of the GB11-Sialyl Lewis A complex was solved using the refined structure of free GB11 as a search model. Structures were refined using Refmac5 [47] followed by iterative model-building cycles using Coot [48]. Restraints for sugars were created using Privateer [49]. Once refinement statistics converged, PDBredo with paired refinement [50] was used to determine whether data of higher resolution shells should be used. The resulting model was once more refined using Coot and Refmac5 as described previously. The relevant data collection and refinement statistics can be found in Table S1. Figures were created using CCP4mg.

### In silico Methods

#### Energy Minimisation and Molecular Docking

Computational studies were carried out to analyse the structural models of mAb-sLeA ligand complexes for GB11 and the well-characterised 1116-NS-19-9 as a reference. Crystallographic structures were available for both mAbs in their apo and holo forms (GB11: apo 9I6Q, holo 9I9H; 1116-NS-19-9: apo 6XUD, holo 6XTG). While these structures provided the basis for modelling, the moderate resolution of the GB11-sLeA complex (3.02–2.80 Å, PDB ID: 9I9H) necessitated additional refinement through molecular modelling techniques, including geometry optimisation and ligand docking. To evaluate the accuracy of the docking protocol, the sLeA ligand was redocked into the apo 1116-NS-19-9 antibody for comparison. The initial heavy-atom coordinates were obtained from the crystallographic structures mentioned above. Missing hydrogen atoms were added using PyMOL [51].

In case of the GB11-sLeA complex, the ligand coordinates were extracted separately from the mAb structure. Both ligands and mAb were solvated in explicit TIP3P water [52] within cubic boxes of 107 x 107 x 107 A^3^ (for GB11) and 39 x 39 x 39 A^3^ (for sLeA). The systems were then neutralised with Na⁺ and Cl⁻ counterions, and CHARMM-GUI tools were used for solvation and ion placement [53]. To optimise the structures of the solvated sLeA and apo GB11 models, energy minimisation was performed using the conjugate gradient algorithm [54,55] and the CHARMM36 force field [56]. Minimisation included 10,000 steps for GB11 and 5,000 steps for sLeA executed in NAMD (Nanoscale Molecular Dynamics, version 2.13) [57,58]. The sLeA tetrasaccharide ligand was modelled as a chain of four monosaccharides: GlcNAc, Gal, Fuc, and Neu5Ac. For non-bonded interactions, a cutoff distance of 12.0 Å was applied, with a smoothing function spanning 10–16 Å. Pair-bonded atoms were excluded from non-bonded interactions, and the atom pair list for non-covalent interaction evaluations was updated every 20 steps.

Structural models of the holo GB11 and holo 1116-NS-19-9 mAbs were generated using rigid docking algorithms in AutoDock Vina [59,60], version 1.5.7. For holo GB11, docking was performed using the energy-minimised coordinates of isolated sLeA and apo GB11, while for holo 1116-NS-19-9, the ligand and mAb coordinates were directly obtained from the crystal structure. During this process, the mAbs were kept rigid, while the sLeA ligand remained flexible, allowing it to adapt to the conformation of the binding cavity. AutoDock tools (ADT) were used for preparing both mAbs and ligand files for docking. The grid box for 1116-NS-19-9 model was constructed around the ligand binding site residues, involving Asp31, Trp33, Asn53, Gly55, Asn56, Arg101 and Ala103 of the heavy chain (H) and Tyr32, Tyr49, Arg50, Tyr91 and Arg96 of the light chain (L). The dimension of the grid box in this model was set to 60 x 60 x 60 Å^3^ (xyz) with a grid space of 0.375 Å. In the case of the GB11 model, the grid box was built around the key residues (Asp31, Trp33, Glu50, Asn53, Ala55, Ile56, Arg101 and Ala103) of the heavy chain and the Tyr32, Tyr49, Arg50, Tyr91 and Arg96 residues of the light chain. The grid box size of GB11 was set to 64 x 64 x 64 Å^3^. Setting the grid boxes as described ensured that all sequence deviations among the binding site residues (K54Y, G55A and N56I) were included. Binding modes (poses) were ranked based on AutoDock Vina’s binding affinity score. The highest-ranked pose was carefully inspected and selected as the starting structure for MD simulations.

A root mean square deviation (RMSD) of less than 0.01 Å between the heavy-atom coordinates of the crystal structure of the 1116-NS-19-9–sLeA complex and those obtained from the redocked model confirmed the reliability of the docking protocol and the predicted affinity scores.

#### Classical Molecular Dynamics Simulation

As a next step, extensive molecular dynamics (MD) simulations were performed [61]. To this end, the top ranked, stable docked sLeA complexes of 1116-NS-19-9 and GB11 were imported into NAMD version 2.13 [57,58]. Additionally, the apo forms of 1116-NS-19-9 and GB11 were simulated as reference systems to identify structural changes induced by ligand binding.

The MD protocol comprised of three major steps: energy minimisation, thermal equilibration, and production run. The CHARMM-GUI solution builder module was used for building the system with the rectangular box type and cubic crystal type of size 112 x 112 x 112 A^3^ for 1116-NS-19-9 and 109 x 109 x 109 A^3^ for GB11 [53]. The initial energy minimisation step consisted of 10,000 steps applying the same parameters as described above. During thermal equilibration, the systems were gradually heated from 0 K to 300 K using the NVT ensemble over 250 ps. A Langevin thermostat with a small friction constant of 1 ps^-1^ was applied to control and maintain a constant temperature [62]. During this phase positional restraints were applied to the protein backbone to prevent structural distortions while allowing solvent and counterions to equilibrate. Next, the equilibration was continued with the production run under NPT ensemble for 200 ns maintaining a pressure of 1.013 bar using Nose-Hoover Langevin piston barostat with an oscillation period of 50 fs and an oscillation decay time of 25 fs [63,64]. To improve statistics of MD trajectory analyses, three replicates of 200 ns MD simulations were performed for the 1116-NS-19-9-sLeA and GB11-sLeA complexes, starting at randomly varied initial velocities.

### Saturation Transfer Difference Nuclear Magnetic Resonance Spectroscopy

After purification, GB11 and HA8 were transferred to a D_2_O-based PBS buffer (pH 7.4) through repeated ultrafiltration employing Amicon Ultra-4 devices (MWCO 30 kDa, Merck Millipore; Tullagreen, Ireland) until the H_2_O-based buffer was sufficiently replaced (under 0.1% residual water). The lyophilised, commercial 1116-NS-19-9 mAb was resuspended in D_2_O-based PBS buffer and trehalose, which was an additive present in the mAb sample, was removed via ultrafiltration. Therefore, the number of ultrafiltration steps for 1116-NS-19-9 was significantly higher than for GB11 and HA8. The 3’-sialyl Lewis A (BIOSYNTH Carbosynth, OS00745) was added from a 20 mM stock solution in the same D_2_O-based PBS buffer (pH 7.4) to a final concentration of 200 µM. The mAb concentration was set to 6.7 µM. The resulting sample was transferred to a 3 mm NMR tube and measured immediately. Negative control samples without mAb were prepared identically, replacing the mAb solution with D_2_O-based PBS (pH 7.4). The sample for the resonance assignment of sLeA was prepared by dissolving the compound in D_2_O- based PBS buffer (pH 7.4) to a concentration of 13.7 mM.

NMR measurements were performed on a Bruker Avance III 600-MHz spectrometer equipped with an N_2_-cooled cryogenic 5-mm TCI-H/C/N-triple resonance probe at 300 K. Resonances of sLeA were assigned from a suite of 2D correlation NMR experiments, namely ^1^H-^13^C-HSQC, ^1^H- ^13^C-HMBC, ^1^H-^1^H-TOCSY, ^1^H-^1^H-COSY and ^1^H-^1^H-NOESY. 1D saturation transfer difference (STD) NMR experiments were acquired using the standard pulse sequence “stddiffesgp.3” from the Bruker library [65]. Water suppression was achieved by excitation sculpting using an 8 ms square pulse [66]. On-resonance saturation of the proteins was achieved by a train of low power Gaussian-shaped pulses at 8.25 ppm with a total saturation time of 2 s. The off-resonance frequency was set to 30 ppm. On- and off-resonance spectra were acquired in an interleaved manner to minimise subtraction artefacts, of which 23552 transients were collected. The on-resonance spectrum was subsequently subtracted from the off-resonance spectrum to yield the difference spectrum that was analysed and quantified. All spectra were acquired, processed and analysed with the software TopSpin (Bruker; v3.6: acquisition; v4.1: processing and analysis). The intensity of the STD effect was quantified by scaling down the off-resonance spectrum to match the intensity of the corresponding signal in the difference spectrum. All measurements were performed in triplicates to allow for statistical analysis. The amounts of the commercial 1116- NS-19-9 needed to carry out triplicates for this assay were beyond our financial means. Therefore, the same sample of 1116-NS-19-9 was measured three times.

## Results

### The New Antibodies Share A High Sequence Similarity

Following the immunisation with synthetic sLeA conjugated to CRM_197_ (Figure S1), we obtained two novel antibodies, GB11 and HA8, using hybridoma technology. RNA isolation followed by RT-PCR was used to obtain DNA to determine the sequences of both mAbs. The light chains of GB11 and HA8 were identified as κ-class (Figure S2). Comparison of the variable regions of heavy and light chain sequences with those of 1116-NS-19-9 (PDB ID: 6XTG) revealed remarkable identities of the variable regions: 97% for GB11 and 96% for HA8 in the variable region of light chains, and 91% for GB11 and 89% for HA8 in the heavy chain sequences (Figure 1a-b). The sole differences in the binding regions are identified in the complementarity-determining region 2 (CDR2) in the heavy chains for both GB11 and HA8. Instead of the charged Lys, and a polar Asn, GB11 features the hydrophobic Tyr and Ile at positions 54 and 56 (sequential numbering of amino acids), respectively. Likewise, HA8 displays a more hydrophobic CDR2, with Trp and Val instead of Lys and Gly at positions 54 and 55, respectively. The "conserved region" of GB11 (PDB ID: 9I9H) reveals several mutations in the CH1 and CL regions, leading to a reduced sequence identity of the heavy chain to 78% and of the light chain to 79% compared to 1116-NS-19-9 (Figure S3).

**Figure 1.**
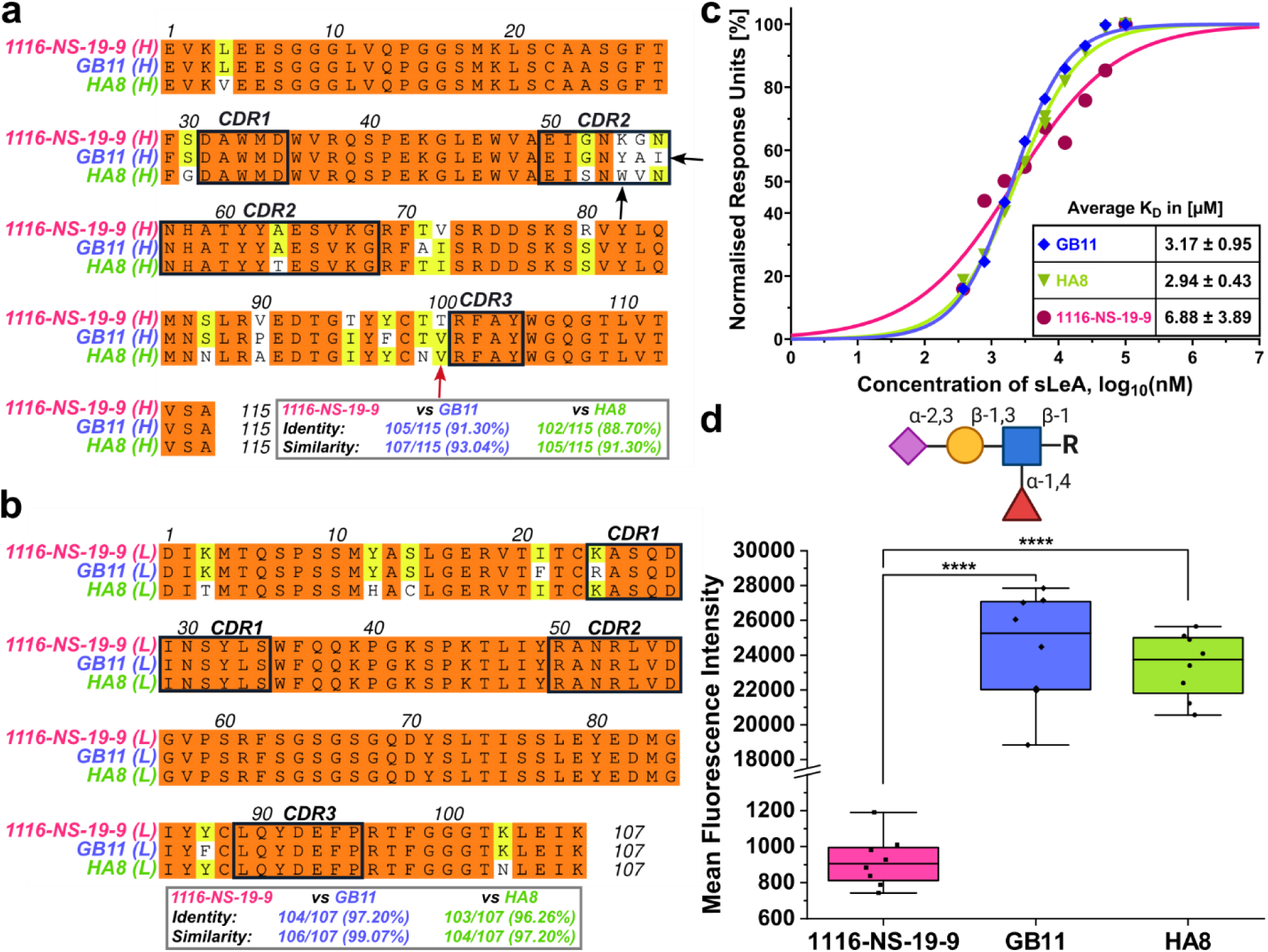
Enhanced binding of GB11 and HA8 to synthetic sLeA despite sequence similarity to 1116-NS-19-9. (**a-b**) Amino acid sequence alignment of heavy (H) and light (L) chain variable regions of 1116-NS-19-9, and the novel mAbs GB11 and HA8. Complementarity-determining regions (CDRs)are indicated by black boxes. Amino acid variations are indicated in white, with arrows highlighting potentially significant mutations. The red arrow signifies one of seven affinity-enhancing key mutations previously introduced by Borenstein-Katz et al. [26]. Identity and similarity scores relative to 1116-NS-19-9 are shown below. (**c**) SPR analysis showing binding of synthetic sLeA to captured mAbs 1116- NS-19-9 (magenta), GB11 (blue), and HA8 (green). One representative run per mAb is shown. Response units (RU) were normalised an plotted against the logarithmic sLeA concentration. Data were fitted using a variable-slope, dose-response model in GraphPad Prism (v10.4.2). KD averages across three replicated are listed in the inset table. (**d**) Mean fluorescence intensity (MFI) of 1116-NS-19-9 (magenta), GB11 (blue) and HA8 (green) binding to synthetic sLeA, immobilised on a glycan array slide. Each mAb was tested in four independent binding assays. Data are presented as box plots; statistical analysis employed one-way ANOVA (**** P ≤ 0.0001). The glycan structure of sialyl Lewis A is depicted in “Symbol Nomenclature for Glycans” (SNFG) notation above (created with BioRender.com).

All three mAbs demonstrate comparable behaviour in a thermal stability assay (Figure S4). HA8, GB11, and 1116-NS-19-9 all display relatively high melting points. HA8 has a melting point of 88°C, while GB11’s melting point is 90°C. Interestingly, 1116-NS-19-9 showed two melting points, with T1 at 79°C and T2 at 91°C. A shoulder in a similar temperature range was also observed for GB11 and HA8 (Figure S4); whether this corresponds to a real second melting point and, thereby, implies a biphasic behaviour could not be determined conclusively.

### GB11 and HA8 mAbs Reveal Enhanced Binding Affinity for Synthetic sLeA

To evaluate the binding affinities of the generated antibodies against the synthetic immunisation target, we employed both surface plasmon resonance (SPR) and isothermal titration calorimetry (ITC). Both methods independently determined dissociation constants (K_D_) in the micromolar (µM) range, consistent with typical affinities observed for anti-glycan Abs [67].

SPR analysis conducted in triplicates yielded K_D_ values ranging from 2.22-4.12 µM for GB11, 2.51-3.37 µM for HA8, and 2.99-10.77 µM for 1116-NS-19-9 (Figure 1c, Figures S6-S7). GB11 and HA8 thus exhibiting slightly higher affinities than 1116-NS-19-9 (table inset in Figure 1c). Comparable K_D_ values were also obtained for GB11 and HA8 via ITC analyses.

The number of binding sites (N) determined with ITC was approximately 2 for both GB11 and HA8 (Table 2), consistent with bivalent binding, where each antigen-binding fragment (Fab) binds the antigen independently. Moreover, the enthalpic changes (ΔH) were similar between GB11 and HA8, indicating that the binding to sLeA likely occurs in a similar manner. Representative titration and fitted enthalpy curves are presented in Figure S5a-c.

**Table 2).**
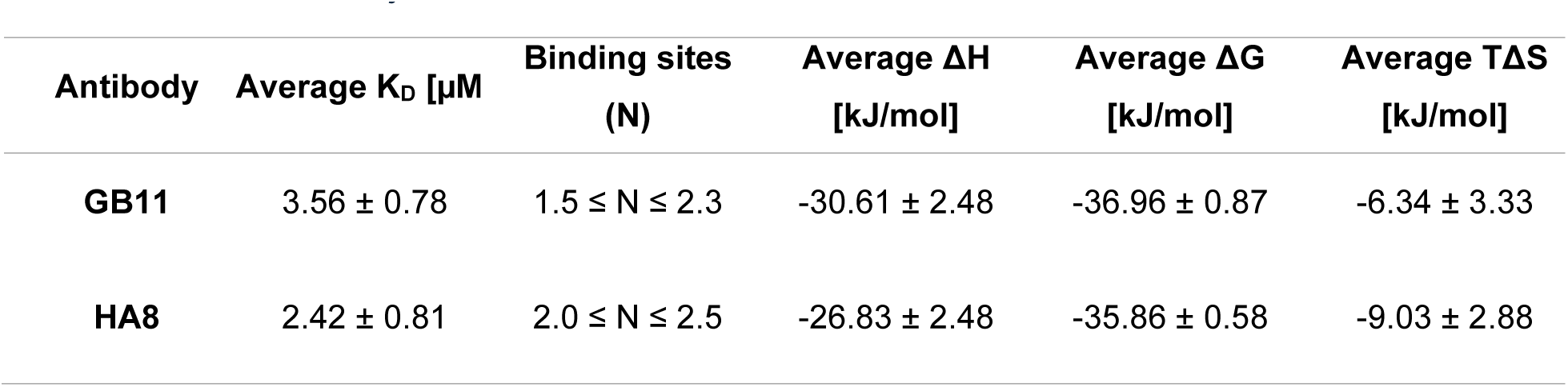
Affinity measurements of GB11 and HA8 via ITC and SPR are congruent. The average values of three independent replicates of GB11 and HA8 in isothermal titration calorimetry analysis. The mAb was in the cell, while sLeA was injected.

| Antibody | Average $K_D$ [$\mu\text{M}$] | Binding sites (N) | Average $\Delta H$ [kJ/mol] | Average $\Delta G$ [kJ/mol] | Average $T\Delta S$ [kJ/mol] |
| --- | --- | --- | --- | --- | --- |
| **GB11** | $3.56 \pm 0.78$ | $1.5 \leq N \leq 2.3$ | $-30.61 \pm 2.48$ | $-36.96 \pm 0.87$ | $-6.34 \pm 3.33$ |
| **HA8** | $2.42 \pm 0.81$ | $2.0 \leq N \leq 2.5$ | $-26.83 \pm 2.48$ | $-35.86 \pm 0.58$ | $-9.03 \pm 2.88$ |

To elucidate antigen-binding specificity of the mAbs, we used synthetic glycan arrays. All three mAbs, GB11, HA8, and 1116-NS-19-9, bound exclusively to sLeA, with no detectable interacting with structurally related glycans, including those of other sialylated Lewis family members (Table 1, Figure S8a, S8e), confirming their high epitope specificity (Figure 1d, and S8b-d). Moreover, GB11 and HA8 exhibited significantly higher mean fluorescence intensities (MFI) than 1116-NS-19-9, suggesting enhanced binding to the immobilised, synthetic target antigen sLeA (Figure 1d). Specifically, HA8 demonstrated a 26.6-fold increase (mean MFI: 24,428.5 ± 3,190.1) and GB11 a 25.5-fold increase (mean MFI: 23,413.9 ± 1,867.5), relative to 1116-NS-19-9 (mean MFI: 916.6± 142.6).

### GB11 Demonstrates Better Binding to Native sLeA Compared to Other mAbs

GB11, HA8, 1116-NS-19-9 bound native sLeA on patient-derived carcinoma tissue of pancreatic ductal adenocarcinoma (PDAC) and gastric adenocarcinoma (GAC) with no apparent differences and minimal binding to cancer-free pancreatic and gastric mucosa tissue (Figure 2a-b).

**Figure 2.**
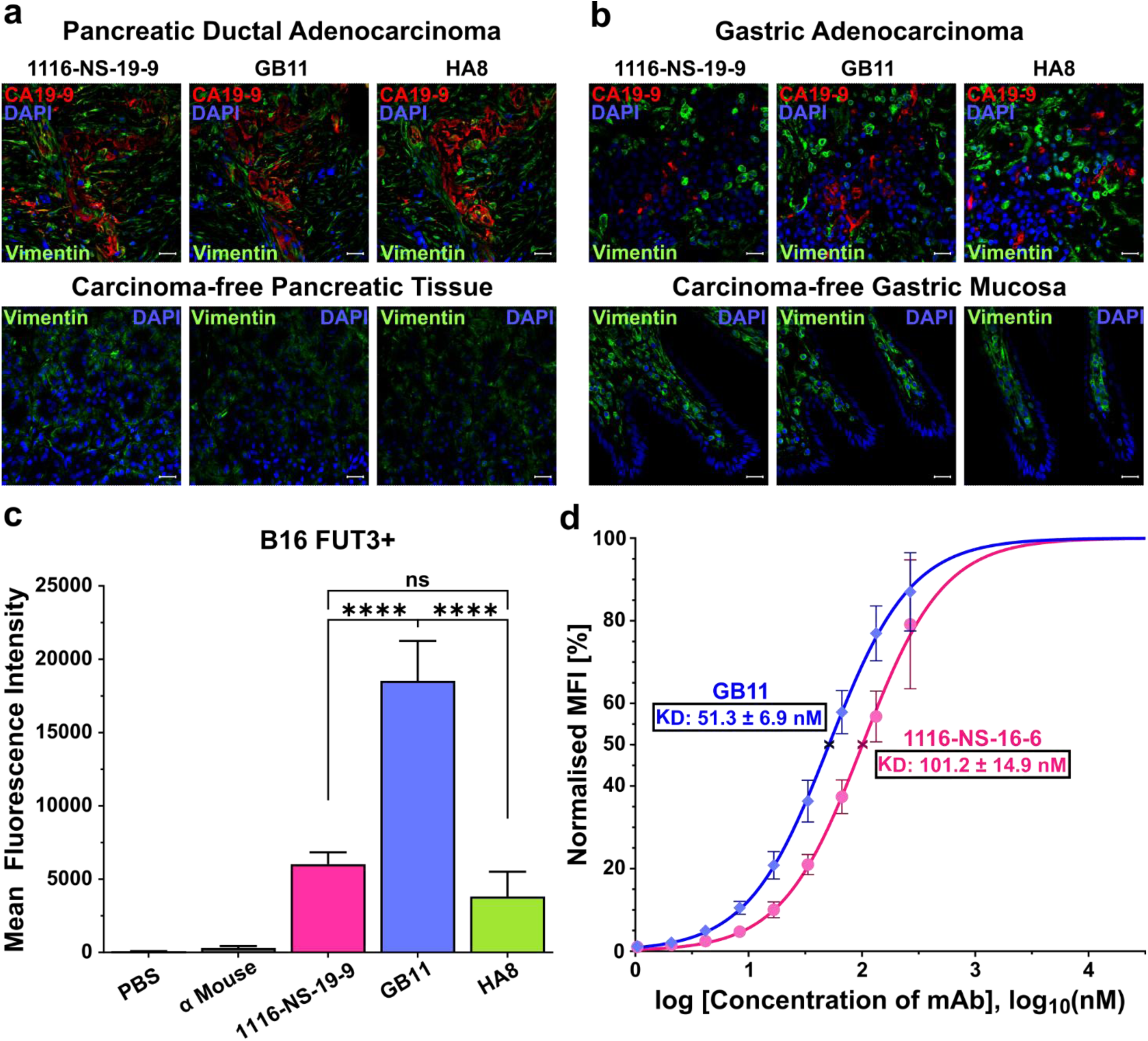
GB11 exhibits significantly stronger binding affinity to native sLeA. (**a-b**) Immunohistochemistry (IHC) revealed binding of α-sLeA mAbs as indicated with sLeA (red), DAPI-stained cell nuclei (blue) and Vimentin (green). The scale bar denotes 20 μm. (**a**) Upper: IHC of pancreatic ductal adenocarcinoma; Lower: carcinoma-free pancreatic tissue. (**b**) Upper: IHC of gastric adenocarcinoma; bottom: carcinoma-free gastric mucosa. (**c-d**) Flow cytometry analysis on sLeA-expressing B16-FUT3+ mouse melanoma cell line reveals (**c**) MFI with SEM of 1116-NS-19-9 (magenta), GB11 (blue) and HA8 (green), with five independent assays per sample. Statistical analysis utilised one-way ANOVA (ns = not significant, **** P ≤ 0.0001). Additionally, a titration binding assay (**d**) for GB11 (blue) and 1116-NS-19-9 (magenta) was conducted, with MFI normalised relative to the highest measured MFI of each mAb. The dots represent the normalised MFI with its corresponding SEM. The fitted results illustrate the normalised MFI against an increasing concentration of each mAb over a logarithmic scale, with the K_D_ values marked with **x** on the fitted curve.

To further evaluate the binding of GB11, HA8, and 1116-NS-19-9 to the native epitope, flow cytometry measurements were conducted using the genetically engineered B16-FUT3+ cell line, which expresses the human fucosyltransferase III (FUT3), resulting in stable sLeA expression on the cell surface [41]. The parental B16 cell line, which lacks sLeA expression, served as a negative control.

All three mAbs bound B16-FUT3+ cells (Figure 2c) but not the parental B16 line (Figure S9a), confirming their specificity for sLeA. Among them, GB11 turned out to be the best binder. It demonstrated the strongest signal in flow cytometry, with a significant 3-fold increase in MFI compared to 1116-NS-19-9 (Figure 2c), supporting the previous data for binding to synthetic glycan. Interestingly, despite HA8 showing a similar binding profile to GB11 for synthetic sLeA, its cellular binding to native sLeA was weaker. GB11 and 1116-NS-19-9 bound 4.8-fold and 1.6- fold more strongly, respectively. Given GB11’s superior performance, GB11 and 1116-NS-19-9 were selected for further evaluation in a flow cytometry-based titration binding assay using B16- FUT3+ cells. This assay yielded apparent K_D_ values of 51± 7 nM for GB11 and 101± 15 nM for 1116-NS-19-9 (Figure 2d, Figure S9b).

### Pre-Organisation of GB11’s Antigen Binding Site Might Lead to Higher sLeA Affinity

Given the relatively minor sequence differences of GB11 and 1116-NS-19-9, yet the significantly higher binding properties of GB11 to synthetic and native sLeA, we sought to better understand their structural divergence using X-ray crystallography. Structures of GB11 were successfully determined in both apo (PDB ID: 9I6Q) and antigen-bound (PDB ID: 9I9H) forms, while attempts to obtain crystals for HA8 were unsuccessful.

The GB11 Fab overlaps with 1116-NS-19-9 with a RMSD of 0.44 and 0.47 Å for the native and the ligand-bound structures, respectively. The sLeA antigen binds within a groove formed by all CDRs except for CDRL1 (Figure 3a). Despite the clearly stronger binding to sLeA by GB11 compared to 1116-NS-19-9, the 3D structures of both complexes are remarkably similar. The sLeA antigen adopts the same low-energy conformation in both structures while interacting with the same amino acids. For GB11 this interaction consists of a hydrogen bond (H-bond) network with Trp33, Asn53, Arg101 of the heavy chain, and Tyr91 and Arg96 of the light chain, along with a single salt bridge between Arg50 of the light chain and the carboxylate of the Neu5Ac (Figure 3a).

**Figure 3.**
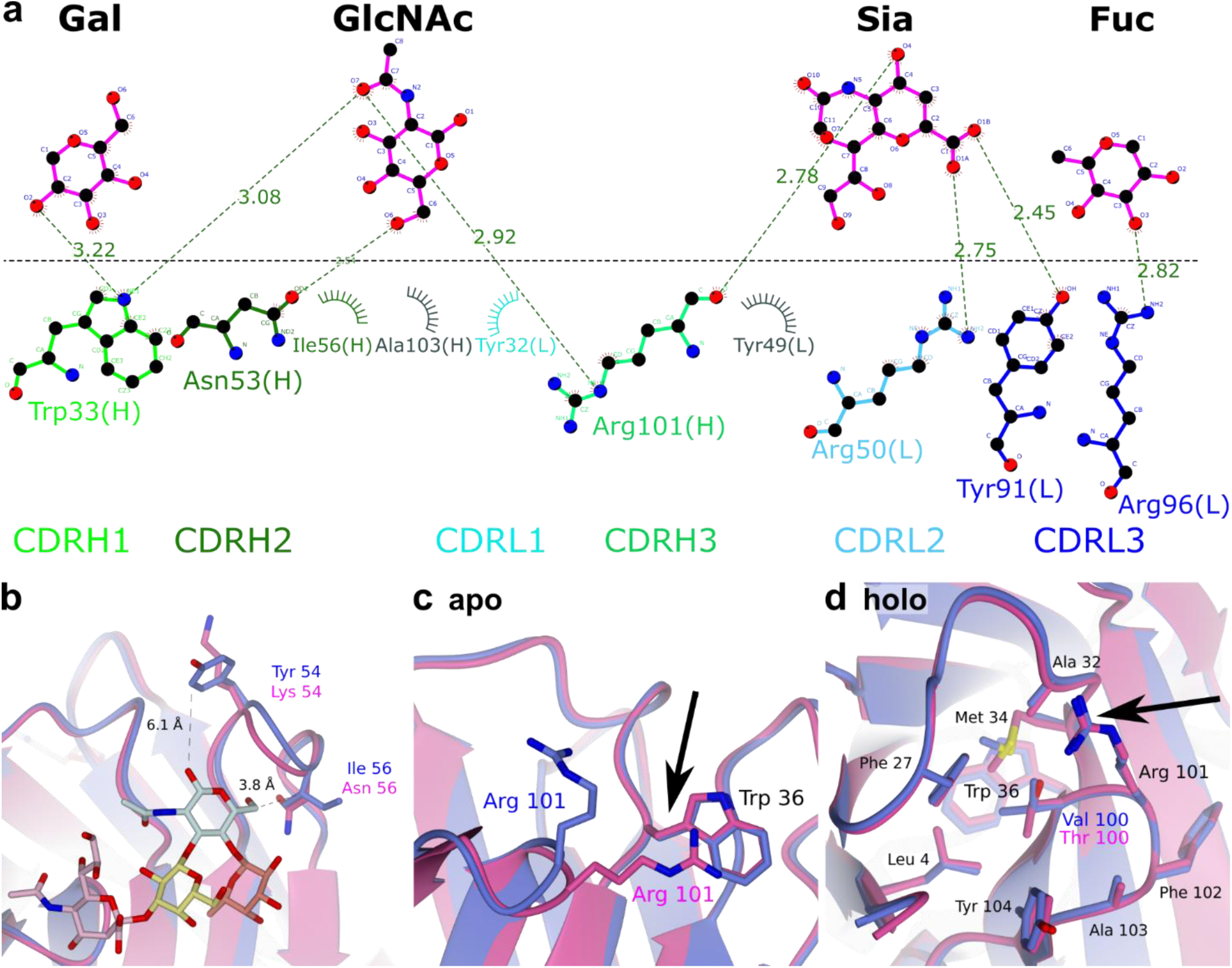
GB11 possesses an open rotamer configuration unlike 1116-NS-19-9, which requires conformational change upon sLeA binding. (**a**) Interaction network between amino acid residues of GB11 and sLeA. (**b**) Overlay of the holo-structure of GB11 (PDB ID: 9I9H; blue) and 1116-NS-19-9 (PDB ID: 6XTG; magenta), displaying side chains of mutations within the CDR of GB11 and their interatomic distances to the antigen sLeA. (**c-d**) Arg101 of 1116-NS-19-9 must undergo a conformational change upon binding, whereas GB11 is already in the binding conformation. (**c**) Overlay between the apo-structure of GB11 (PDB ID: 9I6Q; blue) and 1116-NS-19-9 (PDB ID: 6XUD; magenta). Arg101 in GB11 already adopts an open conformation, whereas in 1116-NS-19-9 it interacts with the nearby Trp36. (**d**) In the holo-structures Arg101 of GB11 and 1116-NS-19-9 are in the same conformation, as indicated by the black arrow, within the hydrophobic pocket created by Val100 for GB11 and Thr100 for 1116-NS-19-9 as well as Leu4, Phe27, Ala32, Met34, Phe102, Ala103 and Tyr104.

Among the differing amino acids within the CDRH1, Tyr54 of GB11 appears to be positioned too distant from the antigen to participate, with a distance of 6.1 Å. Conversely, Ile56 is located at a distance of 3.8 Å from the C6 of the GlcNAc, suggesting a potential involvement in a hydrophobic interaction (Figure 3b). The corresponding Asn56 (Asn53 Kabat numbering scheme [26]) in 1116- NS-19-9 creates a slightly more hydrophilic environment, but with a distance of 3.9 Å for the NH_2_ group and the geometry, it is unlikely to be involved in a direct H-bond.

However, the Thr100Val substitution within the variable heavy chain may play a more significant role in the binding to sLeA. This is supported by the fact that the Arg101 residue within the same loop forms a hydrogen bridge with both the GlcNAc and the Neu5Ac. The hydrophobic pocket, which the side chain of residue 100 must enter for the binding, may be stabilised by the higher hydrophobicity of Val (in GB11) over Thr (in 1116-NS-19-9). Indeed, while the 1116-NS-19-9 antibody’s apo-structure is reported to form a cation-π interaction between Trp36(H) and Arg101(H) (Trp33 and Arg95 Kabat numbering scheme [26]), and possibly also an H-bond between Glu50(H) and Arg101(H), these interactions appear to be absent in our GB11 antibody, with the Arg101 already displaying the same rotamer configuration as in the holo-structure of 1116-NS-19-9 (Figure 3c-d). Thus, the antigen binding site in GB11 is already pre-organised to bind the antigen.

### Molecular Dynamics Simulation Supports Better sLeA Binding Affinity of GB11

To gain deeper insight into carbohydrate recognition and link the high-resolution, yet static details of the X-ray structure with the dynamic situation in solution as reflected by our STD NMR studies (see below), we performed extensive MD simulations.

To maximise quality and information content, we applied an optimised protocol for simulating and analysing protein ligand complexes consisting of energy minimisation, re-docking, and thermal equilibration prior to a long (200 ns) MD production run. For reference purposes, the apo structures of both antibodies were also subjected to MD simulations under identical conditions.

As both mAbs share a high sequence similarity, we expected both models to entertain similar H- bond interaction networks. The 2D LigPlots in Figure S10, generated from the top-ranked tetrasaccharide poses of 1116-NS-19-9 and GB11, illustrate key interactions, with predicted binding affinities of -7.9 kcal/mol and -7.6 kcal/mol, respectively [68]. In both complexes, as detailed in the LigPlot analysis [69], sLeA forms H-bonds with key residues, including Arg101(H) (heavy chain), Arg53(L), and Arg96(L) (light chain). Additional interactions include H-bonds with Trp33(H), Asn53(H), Arg50(L), and Tyr91(L) for sLeA-1116-NS-19-9, and with Glu50(H) and Tyr32(L) for sLeA-GB11.

To investigate the differences in binding site residues between the two antibodies observed in the docking results and determine whether these interactions with sLeA are transient or contribute to stronger binding, we analysed the last 100 ns of all three replicates from the 200 ns MD simulations for each antibody. This analysis provided insights into the stability and compactness of the sLeA-Ab-complexes by evaluating key metrics such as RMSD and Root-Mean-Square-Fluctuation (RMSF) of heavy atoms as well as the Radius of Gyration (Rg), Molecular Mechanics Poisson-Boltzmann Surface Area (MM-PBSA), and interaction energy scores.

The time courses of the backbone RMSD scores relative to the starting structure reveal that mAb 1116-NS-19-9 converges to a stable bound conformation within the final 50 ns of the simulation, with an average RMSD deviation of less than 2 Å. For GB11, convergence is achieved within the last 20 ns, with an average RMSD deviation of less than 1.5 Å in two replicates and less than 2 Å in one replicate. Notably, in both complexes, sLeA converges from its initial structure (generated by redocking) to a highly stable bound conformation, with average RMSD deviations of less than 1 Å (Figure S11a-d).

The mobility of individual residues during the simulation was assessed using RMSF. We focused on RMSF deviations within the binding site residues of both antibodies, comparing their apo and holo forms to evaluate the impact of sLeA binding. Notably, sLeA binding to GB11 reduced the motional amplitude of binding site residues by up to 1 Å (Figure S12b), whereas no such reduction was observed in 1116-NS-19-9 (Figure S12a). This decrease in flexibility, indicative of increased backbone rigidity, suggests that GB11 forms a more stable binding interface with sLeA.

As a measure for quantifying the compactness and stability of biopolymers and biopolymer complexes, we calculated the radius of gyration (Rg) [70]. The plots reveal that the mAb structures maintain a high degree of compactness, with deviations of Rg values relative to the start conformations within the permissible limit of less than 1 Å in both the apo and holo forms of 1116-NS-19-9 and GB11. This is further evidence for the stability of the antibodies and their complexes throughout the simulation period (Figure S13).

Notably, we found important support for stronger binding of the sLeA glycan to GB11 by analysing the MD trajectories. To this end, we estimated the enthalpic contributions to the binding free energies of the holo complexes according to their molecular mechanics with Poisson–Boltzmann and surface area solvation (MM-PBSA) scores over the last 10 ns of the simulations [71,72]. The average MM-PBSA of the 1116-NS-19-9 complex was -3.5 kcal/mol, while that of the GB11 complex was -5.2 kcal/mol, indicating a slightly higher affinity of GB11 towards sLeA. This is also reflected in the interaction energies, considering electrostatic and Van der Waals interactions between the antibody and sLeA. The average interaction energy score across the replicates of 1116-NS-19-9 is -65.3 kcal/mol, while for GB11 it is -68.4 kcal/mol (ΔΔH ∼ 3.1 kcal/mol). Furthermore, when comparing the time frames representing the lowest interaction energy scores among the replicates of the antibody complexes, the 1116-NS-19-9 complex displayed a score of -81.7 kcal/mol, while the GB11 complex attained a score of -87.6 kcal/mol (Figures S14-15). This amounts to an even higher difference of ∼6 kcal/mol between 1116-NS-19-9 and GB11 in favour of stronger binding to sLeA by GB11.

These global results can be rationalised by analysing the detailed binding modes and the individual interactions within the binding sites of the antibodies. To this end, we identified the residues involved in forming strong H-bond interactions. Recurring H-bonds across all replicates of the simulations are summarised in a bar diagram (Figure 4a) [73,74].

**Figure 4.**
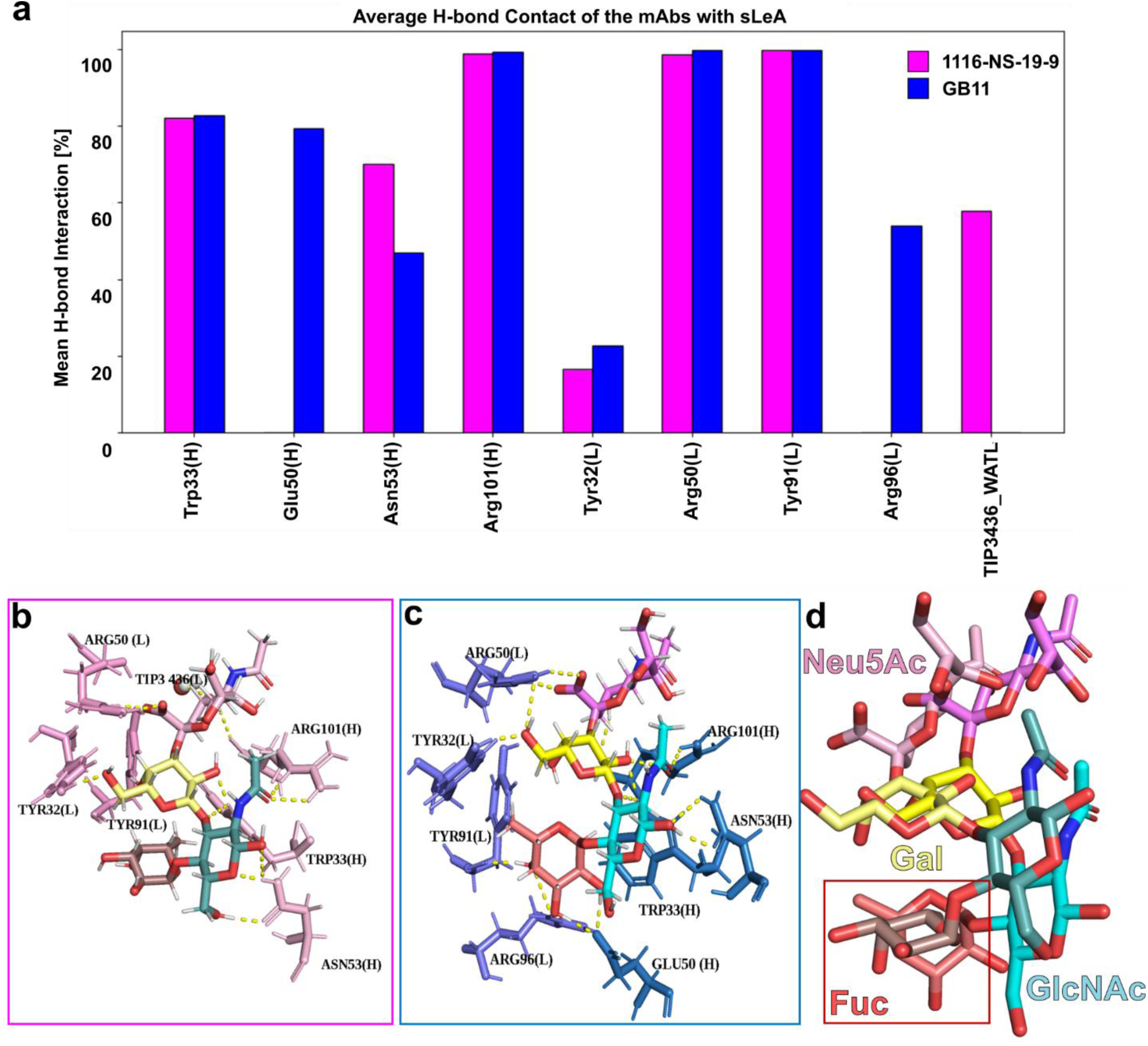
GB11 engages sLeA through distinct H-bonding and a different fucose orientation compared to 1116- NS-19-9. (**a**) Key binding site residues of 1116-NS-19-9 (magenta) and GB11 (blue) involved in forming H-bond interactions with sLeA. The bars represent the stability of each hydrogen bond, shown as the average percentage of simulation time during which the bond was present. (**b-c**) Superimposed binding poses of sLeA at the lowest interaction energy in 1116-NS-19-9 (**b**) and GB11 (**c**). (**d**) In GB11, the Fuc unit adopts a rotated, vertical orientation that facilitates H-bond formation with Glu50(H), Tyr91(L), and Arg96(L). This conformational shift emerges as a key factor contributing to the enhanced binding observed for GB11. In contrast, the horizontal or flat orientation observed in the Fuc, while binding to 1116-NS-19-9, does not promote stable H-bond formations with binding site residues.

The H-bonds maintained throughout the MD simulation period are presented as a percentage with respect to the simulation time. For 1116-NS-19-9, H-bond interactions with sLeA were predominantly observed with the Neu5Ac unit, involving the residues Arg101(H), Arg50(L), and Tyr91(L) for ∼99% of the simulation time, and with the explicit water molecule TIP3436(L) for about 50% of the time. The Gal subunit formed H-bonds with Trp33(H) and Tyr32(L), and the GlcNAc with Asn53(H), while stable H-bonds with the Fuc were not observed.

For GB11, the Fuc formed stable H-bonds with the residues Glu50(H), Tyr91(L), and Arg96(L) for over 90% of the simulation time. The Gal unit mainly formed H-bonds with Trp33(H) and Tyr32(L), and the GlcNAc moiety with Trp33(H) and Asn53(H). Finally, the Neu5Ac unit established H- bonds with Arg101(H) and Arg50(L) for over 90% of the time. In comparison to 1116-NS-19-9, GB11 exhibits stable H-bond interactions with all four carbohydrate residues (Neu5Ac, Gal, GlcNAc, and Fuc) with the binding site residues, with Glu50(H) being particularly specific for H- bond formation with Fuc.

For both mAbs, we took the timeframes exhibiting the lowest interaction energy scores to generate representative snapshots of the sLeA ligand in the binding sites (Figure 4b,4c). It became evident that, in the MD simulation, the sLeA tetrasaccharide as a whole adopts a slightly shifted position in the binding site, and, particularly, the orientation of the Fuc is rotated by ca. 90°. This altered binding mode allows for the Fuc unit to engage more strongly in H-bonds with the key residues of the binding pocket, particularly the Tyr91(L). In contrast, Tyr91(L) of 1116- NS-19-9 forms an H-bond with the monosaccharide unit Neu5Ac instead.

In summary, our analysis of MD simulations reveals the formation of stable mAb-glycan complexes. The slight differences of the binding sites of both mAbs bring about small changes of the binding mode reflected by differential engagement in H-bonds and conformational changes of the sLeA ligand and the binding site residues. This finally results in higher binding affinity of sLeA towards GB11, in line with our experimental results.

### STD NMR Spectroscopy Confirms Unique sLeA Binding Epitopes

To gain deeper insight into the differential recognition of sLeA by the three mAbs, we employed STD NMR spectroscopy. Binding was confirmed in all cases and further supported by negative control experiments conducted in the absence of mAbs (Figure S18).

A significant portion of the tetrasaccharide’s resonances fell within the same ppm range (3.2- 4.2 ppm) and was, therefore, prone to strong overlapping, obstructing an unambiguous quantification of STD effects for specific resonances within this range. Hence, based on reported resonance values for sLeA derivatives [75], we carefully re-assigned the resonances of sLeA under our experimental conditions using conventional two-dimensional NMR techniques (Table S2, Figure S16, S17). Accurate resonance assignment was essential to maximise both the interpretability and reliability of the subsequent STD NMR experiments. Of note, the acquired sLeA sample contained a mixture of α- and β-GlcNAc anomers, attributed to the presence of a free OH-group at the C1 position of GlcNAc. Accordingly, we refer to these forms as the α- and β-anomers, respectively.

While in all three cases, dissociation of the complexes is sufficiently fast to bring about decent sensitivity in STD NMR measurements, 1116-NS-19-9 displays significantly stronger STD effects in terms of the absolute values compared to the other two mAbs (Figure 5a, Figure S19, S20), which is most likely caused by faster off-rates in line with its slightly lower affinity. Therefore, to avoid bias towards a certain NMR signal of sLeA, we normalised the STD effects relative to the sum of all the STD effects separately for every mAb (Figure 5b).

**Figure 5.**
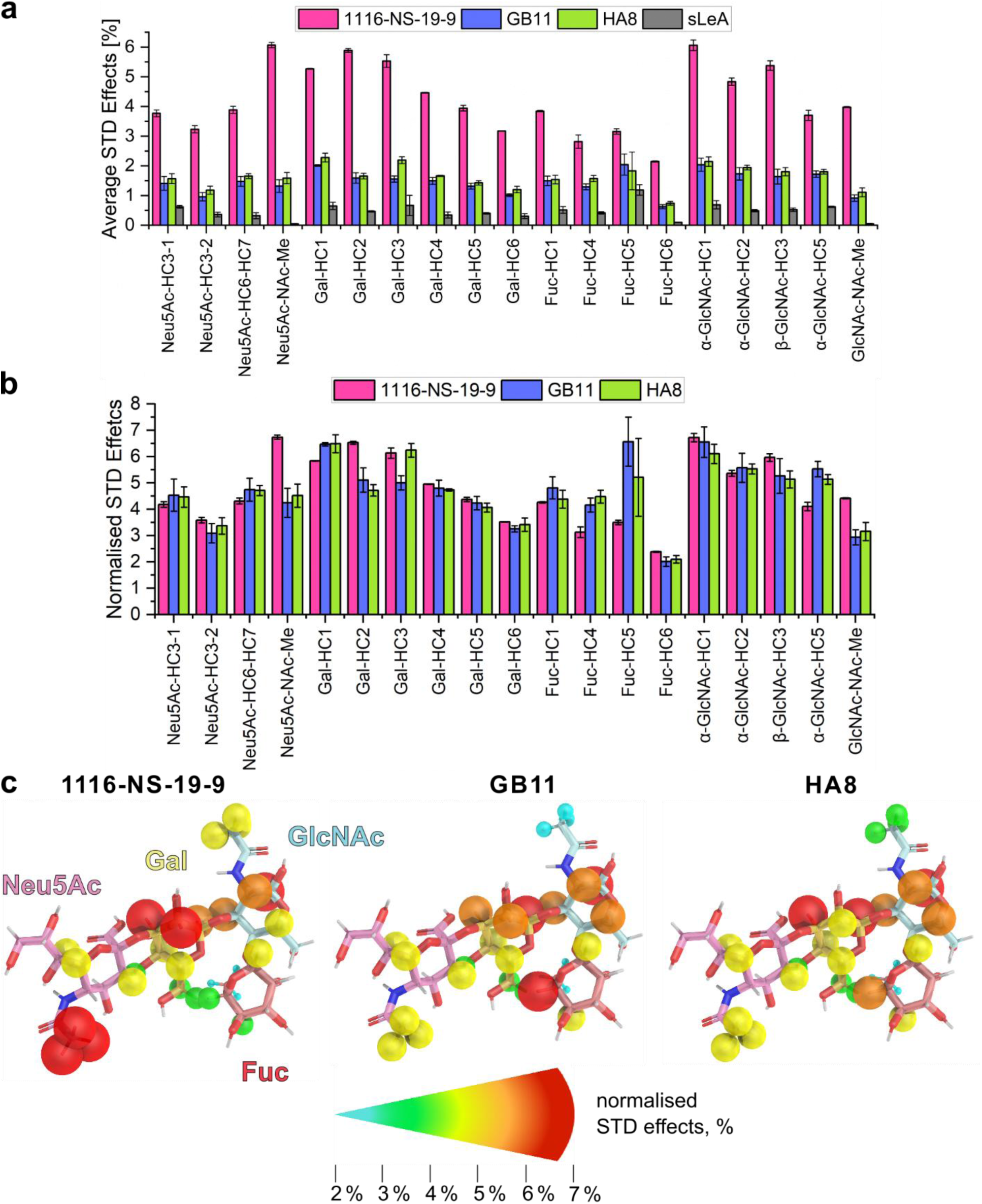
STD NMR reveals distinct sLeA binding epitopes for the three mAbs. (**a**) Average STD effects for sLeA hydrogen atoms from three independent STD NMR measurements with mAbs 1116-NS-19-9 (magenta), GB11 (blue) and HA8 (green). STD effects are expressed as a percentage of the respective sLeA signal without saturation. sLeA (grey bars) serves as a negative control STD NMR experiment in the absence of the mAbs. (**b**) Normalised STD effects relative to the sum of all STD effects for sLeA hydrogen atoms. (**c**) Normalised STD effects are mapped on the molecular structure of sLeA for each mAb, with increasing STD effects are presented by colour and size.

The normalised STD effects mapped on the molecular structure of sLeA depict a rather similar distribution of the STD effects in the entire molecule for GB11 and HA8, with the strongest STD effects concentrated on the Gal, GlcNAc, and Fuc subunits (Figure 5c). In contrast, 1116- NS-19-9 exhibits a different pattern, with the strongest STD effects observed on the Neu5Ac, Gal, and GlcNAc subunits. Interestingly, the acetyl groups of Neu5Ac and GlcNAc feature higher STD effects for 1116-NS-19-9 compared to GB11 and HA8 (Figure 5c).

## Discussion

Cancer patients with advanced and metastatic cancers often suffer from poor prognosis and limited therapeutic options. However, effective mAbs targeting distinct cancer antigens, used for early cancer detection and as mono [76–78], adjuvant [79,80], or drug-conjugate [81,82] therapy, drastically elongate patients’ survival. Our study focused on developing improved mAbs against sLeA, a tumour-associated carbohydrate antigen overexpressed in various cancers with poor survival rates.

To evaluate the effect of the antigen, we decided to use a classic immunisation strategy with a glycoconjugate carrying synthetic sLeA. Two mouse-derived mAbs, GB11 and HA8, were isolated and characterised. These were compared to 1116-NS19-9, an anti-sLeA mAb produced through immunisation against the native, heterogeneous antigen as presented on the human CRC cell line SW1116 [33,34]. A combined biophysical and bioinformatics’ study in comparison with *in situ* data gave a comprehensive overview of the antigen binding process and provided clues for the improved specificity and affinity of GB11, compared to 1116-NS-19-9, despite very high sequence similarity between both mAbs.

The K_D_ values obtained from both ITC and SPR measurements for GB11 and HA8 are consistent with each other (Table 2, Figure 1c), with a binding affinity within the low micromolar range with synthetic sLeA in solution. SPR also shows a similar K_D_ for 1116-NS-19-9 (Figure 1c). Furthermore, all three mAbs exhibit no cross-reactivity towards glycans structurally similar to sLeA in glycan array assays (Figure S8b-d). Comparison of the variable regions of heavy and light chain sequences of GB11 and HA8 with those of 1116-NS-19-9 revealed remarkably high similarities in the variable regions (Figure 1a-b). The variations in the binding affinities of the mAbs, however, may be attributed to the small sequence differences in the CDR2.

Typically, germline Abs are recognised for their inherent polyspecificity due to heightened flexibility in their variable regions, owing to their limited mutational load in the germline V, D, and J genes [83,84]. During the process of affinity maturation, Abs evolve through somatic hypermutations towards greater specificity and affinity for their target antigens [85], often by stabilising the conformation and limiting the flexibility of the antibody’s paratope [86,87].

Carbohydrate antigens, however, present a unique case. Because many tumour-associated glycans share high structural similarity with self-glycans, germline Abs recognising these antigens are often intrinsically selective to avoid autoreactivity [83,88–90]. This pre-existing selectivity may explain why affinity-matured Abs against such targets show relatively minor sequence divergence from their germline precursors. In line with this, the high sequence similarity observed between GB11, HA8, and the earlier mAb 1116-NS-19-9 could reflect a shared origin from germlines genes already predisposed to selectively recognise sLeA. This notion is supported by previous studies showing that germline antibodies targeting tumour-associated glycans can exhibit high selectivity with minimal affinity maturation [88,89]. While our data do not directly confirm this mechanism, the observed similarities are consistent with such a model.

Despite their high sequence similarity, GB11 and HA8 exhibited more than a 25-fold higher MFI than 1116-NS-19-9 in synthetic glycan array assays, suggesting significantly stronger binding to immobilised, synthetic sLeA (Figure 1d). This difference may result from the fact that GB11 and HA8 were generated using a synthetic sLeA glycoconjugate for immunisation, in contrast to 1116- NS-19-9, which was raised against native sLeA displayed on CRC cells. Therefore, we next aimed to investigate whether the observed binding performance would also be mirrored in the binding to sLeA in its native cellular context. Indeed, both GB11 and HA8 identified sLeA also on human PDAC and GAC tissue (Figure 2a-b). Moreover, GB11 demonstrated at least a twofold increase in binding to B16-FUT3+ cells compared to 1116-NS-19-9 (Figure 2d, Figure S9b).

To understand the origin of the observed binding differences as well as to gain insight into the role of sequence variation in the variable regions, we next analysed the crystal structures, MD simulations, and STD NMR data of the mAb–sLeA complexes. The biological role of glycans is closely linked to their structural flexibility, particularly the geometries of glycosidic linkages and the puckering of individual monosaccharide rings [91,92]. These intrinsic dynamics can influence epitope presentation and antibody recognition. Despite their general flexibility [93,94], Lewis antigens like sLeA can adopt relatively rigid “closed” conformations, stabilised by an intramolecular CH···OH hydrogen bond between the Fuc and Gal residues [95–98]. Although conformational transitions within sLeA are not directly captured in our data, transitions to alternative or “open” conformations have been observed in lectin complexes [99] and proposed in computational studies [91,92,100].

In light of these considerations, we next examined how sequence variation and glycan orientation influence the architecture of the binding site. We therefore began by analysing the available crystal structures. Notably, specific binding pocket residues, such as Trp36 and Arg101, exhibited conformational changes upon ligand binding in case of 1116-NS-19-9, as previously reported [26]. Conversely, analysis of the apo- and holo-crystal structures of GB11 revealed that these same amino acids were already configured in the rotamer conformation that is observed in case of 1116- NS-19-9 only upon binding to sLeA and not in the free state (Figure 3c-d). The pre-organisation of GB11 mirrors the binding behaviour observed in the mAb 5B1 [26], which was also generated through immunisation with synthetic sLeA conjugated to KLH [27].

Additionally, we found differences in the binding pocket formed by the heavy chain, where Thr100 of 1116-NS-19-9 was substituted with the more hydrophobic Val100 in GB11 and HA8, potentially contributing to the observed variation in the rotamer conformation of Arg101 by creating a more hydrophobic binding pocket (Figure 3c-d). Indeed, some of the mutations were previously introduced in a directed evolution campaign to improve sLeA binding in 1116-NS-19-9 [26], achieving affinity gains comparable to those measured for GB11 and HA8. Beyond the binding site, specificity can also be modulated through alterations distal to the paratope [101–103]. Indeed, the comparison between GB11, HA8, and 1116-NS-19-9 revealed that mutations predominantly occur outside the core binding site (Figure 1a–b). In case of GB11, some occur even within the structurally conserved region (Figure S3). Such distal mutations may further contribute to the enhanced affinity of GB11 and HA8 by indirectly stabilising the paratope or by modulating the VH-VL domain spacing and orientation, a phenomenon already observed for other anti-glycan Abs [88,89,102,104].

To support our structural studies, comprehensive MD trajectory analyses of the sLeA-mAb complexes in solution phase were performed. In both complexes, the glycans resided stably in the binding pockets after the equilibration phase. Notably, the sLeA-GB11 complex exhibited a slightly lower interaction enthalpy than sLeA bound by 1116-NS-19-9 (1.7 kcal/mol averaged over the last 10 ns). This tendency is corroborated by the predicted interaction energies that are also more favourable in case of the GB11-sLeA complex (3.1 kcal/mol) underpinning the higher affinity of GB11.

The sLeA tetrasaccharide adopts a very similar conformation when binding to the mAbs 1116- NS-19-9 and GB11, and such a conformation is also seen in complex with 5B1 [26]. However, when comparing in detail the conformational space of sLeA docked in 1116-NS-19-9 and in GB11, we observed notable differences in the orientation of the Fuc unit (Figure 4b-d). When superimposing the sLeA tetrasaccharide of both complexes, the Fuc in the GB11 complex adopts a vertical orientation compared to its orientation in the 1116-NS-19-9 complex. This is likely to play an important role in promoting stronger H-bond interactions with the key residues of the binding pocket (Glu50(H), Tyr91(L), and Arg96(L)). Conversely, in 1116-NS-19-9, Tyr91(L) forms an H-bond interaction with Neu5Ac instead. According to our MD simulations, this altered orientation of the Fuc results in stronger H-bond formation and promotes binding contributions of all four glycan units thus further corroborating the overall observed stronger binding affinity of GB11 towards sLeA.

It should be pointed out that these conformational changes and H-bonds are of transient nature and reflect the dynamic situation at room temperature in solution. Noteworthy, also in the X-ray structure, corresponding to an averaged lowest energy conformation in a frozen crystal, the electron density of the Fuc residue binding to GB11 is lowest of all four monosaccharide building blocks. The conformation and position within the binding site are, therefore, less well defined than for the rest of the sLeA ligand and may indicate increased flexibility. That the shifted and rotated orientation of the Fuc moiety may be relevant for GB11’s higher affinity towards sLeA is corroborated by the results of our STD NMR experiments. The altered positioning of the Fuc unit as observed in the MD simulation of the GB11 complex would orient its hydrophobic face, comprising of H3, H4, H5, and the CH_3_-group, closer towards the side chain of Ile-56(H). This would explain the stronger saturation of Fuc-H4 and -H5 in presence of GB11 that is indicative of a hydrophobic interaction that, in turn, is likely to contribute to higher affinity of sLeA towards GB11 compared to 1116-NS-19-9 (Figure 5a-c).

Interestingly, the MD simulations of sLeA in complex with 1116-NS-19-9 reveal a unique H-Bond between Tyr91(L) and the sialic acid moiety Neu5Ac. This is evidence for a stronger contribution of this part of the sLeA-tetrasaccharide that is nicely reflected by increased saturation of the Neu5Ac-acetyl-CH_3_ group in the STD NMR experiments.

When comparing the saturation received by the individual carbohydrate units in the mAb complexes, the STD NMR data for 1116-NS-19-9 show strongest saturation for Gal and GlcNAc, followed by Neu5Ac, with the weakest signal at Fuc (Figure 5, Figure S21). In contrast, GB11 and HAS show more uniform saturation across all four carbohydrate residues (Figure 5, Figure S21), again highlighting stronger engagement of Fuc as observed in the MD simulations.

As nicely depicted in Figure 5a, 1116-NS-19-9 produces higher overall STD signals. Since STD effects are not only dependent on the affinity but also greatly on the dissociation kinetics of the ligand [65,105], it is always the combination of these two factors that determines the absolute value of the STD effects. Considering that SPR provided similar K_D_ values for all three mAbs (Figure 1c), it is reasonable to attribute the differences in STD effects primarily to disparities in dissociation kinetics. In this context, higher STD effects for 1116-NS-19-9 indicate a faster rate of dissociation of sleA compared to GB11 and HAS. The slower binding kinetics observed for the two new mAbs suggest a higher specificity of the binding pockets for GB11 and HAS. These observed kinetic differences further support the binding disparities seen in our experimental results and validate our computational simulations.

The combined insights from X-ray crystallography, STD NMR and MD simulations not only reinforced our experimental results regarding the elevated binding affinity but also provided additional detailed insights into the molecular recognition of each complex, including the key interactions of sleA with each mAb. Moreover, it corroborated the conclusion that GB11 engages sleA more extensively than 1116-NS-19-9. While GB11 interacts with all four monosaccharides units, 1116-NS-19-9 primarily contacts Neu5Ac, Gal, and GlcNAc (Figure 4b-c, Figure 5b-c, Figure S10, Figure S21). Increasing the number of glycan residues in contact with the antibody binding site has been shown to enhance both specificity and affinity by promoting a more extensive interaction network and stabilising a favourable energetic conformation of the glycan [S4]. Hence, GB11 is one example of gaining stronger affinity by enhancing the interaction interface to the carbohydrate antigen. A similar interaction pattern has been described for 5B1, which also engages all four residues [26], suggesting that full epitope recognition is a recurring feature among high-affinity anti-sleA antibodies.

In summary, our comprehensive analysis and previous studies [26,27] suggest that the strategic use of synthetic glycans in immunisation can lead to the development of anti-glycan Abs exhibiting higher binding specificity and affinity towards their target. Furthermore, using novel characterisation of sleA binding by STD-NMR, we unveiled key determinants that shed light on the way these Abs may achieve superior performance through pre-organised binding sites that precisely recognise distinct glycan configurations and engage the carbohydrate ligand through a broader contact surface.

## Conclusion

The sialyl Lewis A antigen, or CA 19-9, is an FDA-approved marker for pancreatic ductal adenocarcinoma (PDAC), an aggressive malignancy with extremely poor survival rates [106]. High-affinity human mAbs targeting sLeA are currently undergoing clinical trials for PDAC therapy [107,108]. Developing novel and improved tools for specifically targeting PDAC, colorectal, and other sLeA-expressing cancers could significantly improve their treatment and patient survival. This work demonstrated the advantages of using well-defined, synthetic glycan antigens for the development of highly selective mAbs against tumour-associated carbohydrate antigens and unveiled the structural aspects behind the improved properties of the newly generated mAbs.

## List of Abbreviations

Ab/Abs: Antibody / antibodies
CA 19-9: Carbohydrate antigen 19-9 (equals sLeA)
CDR: Complementarity-determining region
CRC: Colorectal carcinoma
DAGR: Database of Anti-Glycan Reagents
Fab: Fragment antigen-binding region
FDA: Food and Drug Administration
Fuc: Fucose
FUT3: Fucosyltransferase III
Gal: Galactose
GAC: Gastric adenocarcinoma
GlcNAc: N-Acetylglucosamine
H-bond: Hydrogen bond
IgG: Immunoglobulin G
IHC: Immunohistochemistry
ITC: Isothermal titration calorimetry
K_D_: Dissociation constant
mAb/mAbs: Monoclonal antibody / antibodies
MD: Molecular dynamics
MFI: Mean fluorescence intensity
MM-PBSA: Mechanics Poisson-Boltzmann Surface Area
Neu5Ac: N-Acetylneuraminic acid
NMR: Nuclear magnetic resonance
PDAC: Pancreatic ductal adenocarcinoma
Rg: Radius of gyration
RMSD: Root-mean-square deviation
RMSF: Root-mean-square fluctuation
RT-PCR: Reverse transcriptase-polymerase chain reaction
sLeA: sialyl Lewis A (equals CA 19-9)
SPR: Surface plasmon resonance
STD: Saturation transfer difference
TACA: Tumour-associated carbohydrate antigen WT Wilde type

## Declarations

### Ethics approval and consent to participate

Animal experiments were performed by Hybrotec GmbH (Germany, Potsdam) and approved by the Landesamt fur Arbeitsschutz, Verbraucherschutz und Gesundheit (LAVG) Brandenburg (Gesch-Z. 2347-A-34-1-2020). Experiments were performed according to the German law, following the regulations of the Society for Laboratory Animal Science (ALAS) and of the Federation of Laboratory Animal Science Associations (FELASA).

Patients provided consent for the collection, encryption, storage, and analysis, including genotyping, of their tissue and blood samples for tumour research in a pseudonymised form by the staff of the "Gewebe- und Blutprobenentnahme fur die Tumor- und Biobank (TBB)" at the "Charite Comprehensive Cancer Center" (CCCC) under approval number EA1/152/10. They also consented to the pseudonymised recording and processing of their data on electronic media and agreed to the anonymous publication of research results, ensuring their identity remains undisclosed.

### Consent for publication

Not applicable.

### Availability of data and materials

All presented data and materials are available upon request from the corresponding authors.

### Authors’ contributions

O.M., H.M.M., A.F, and S.K.K. designed the research. F.G., A.F., S.K.K.,J.L. generated and isolated the Monoclonal Antibody. S.K.K. generated, analysed, and interpreted the ITC data. A.F., S.K.K., and G.M.S.G.M. performed, analysed, and interpreted the SPR data. IHC was performed by J.A. Thermal Shift Assay, Glycan Array Assays, and Flow Cytometry were executed, analysed, and interpreted by A.F. Crystallisation has been performed by S.K.K. and C.R. X-Ray analysis was conducted by M.K. Molecular Dynamics Simulation was carried out by S.M.K and analysed by S.M.K. and M.A.M. STD-NMR was conducted and analysed by R.N. and A.F. S.K.K. and A.F. wrote the manuscript with contributions from R.N., S.M.K., M.K., C.R., M.A.M., H.M.M., and O.M.. Supervision was given by C.K., M.A.M., C.R., P.H.S., H.M.M., and O.M. The final manuscript was approved by all authors.

## Acknowledgment

We greatly acknowledge the technical support provided by Katrin Sellrie throughout the hybridoma selection process. We also extend our gratitude to Prof. Dr. J. V. Ravetch for generously supplying the B16 and B16FUT3+ mouse melanoma cell lines. We would like to thank Ken Motohashi for his generous contribution of the pCRZeroT plasmid. Additionally, we appreciate the assistance of Hybrotec GmbH during the immunisation process and GlycoUniverse GmbH for the synthesis of sLeA. C.R. thanks the beamline staff from the HZB Berlin for the support during data collection. A.F. expresses sincere gratitude to colleagues, friends, and family for their ongoing encouragement and assistance during a period of significant health challenges. Special thanks from her side go to S.K.K., R.N., O.M., and H.M.M. for their patience, understanding, and continued support.

