## Supplementary Information for "An Integrative Approach to Develop and Characterise Antibodies Against the Cancer Associated Antigen Sialyl Lewis A (CA 19-9)"

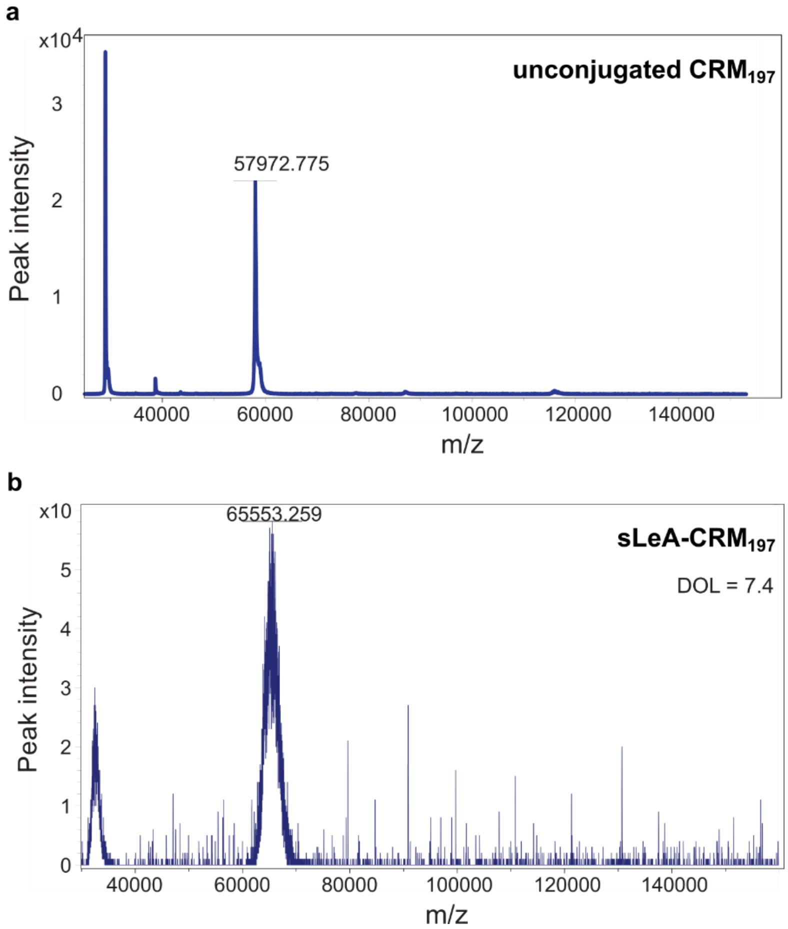


***Figure S1| Successful conjugation of sLeA to CRM$_{197}$.*** *MALDI-TOF spectra of unconjugated CRM$_{197}$ (**a**) and CA 19-9 conjugated to CRM$_{197}$ (**b**), depicting a loading of approximately 7.4 sLeA molecules/protein.*

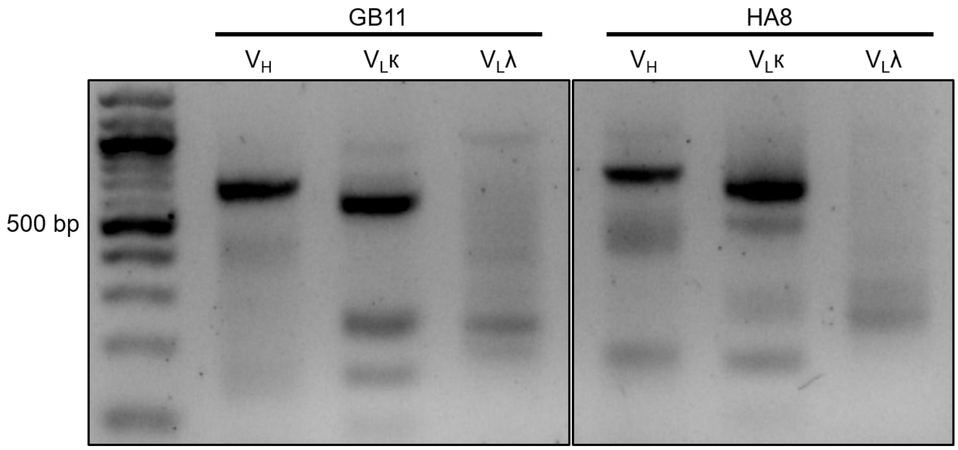


***Figure S2| RNA extraction and RT-PCR, revealing the light chain of GB11 and HA8 is a κ-chain.***

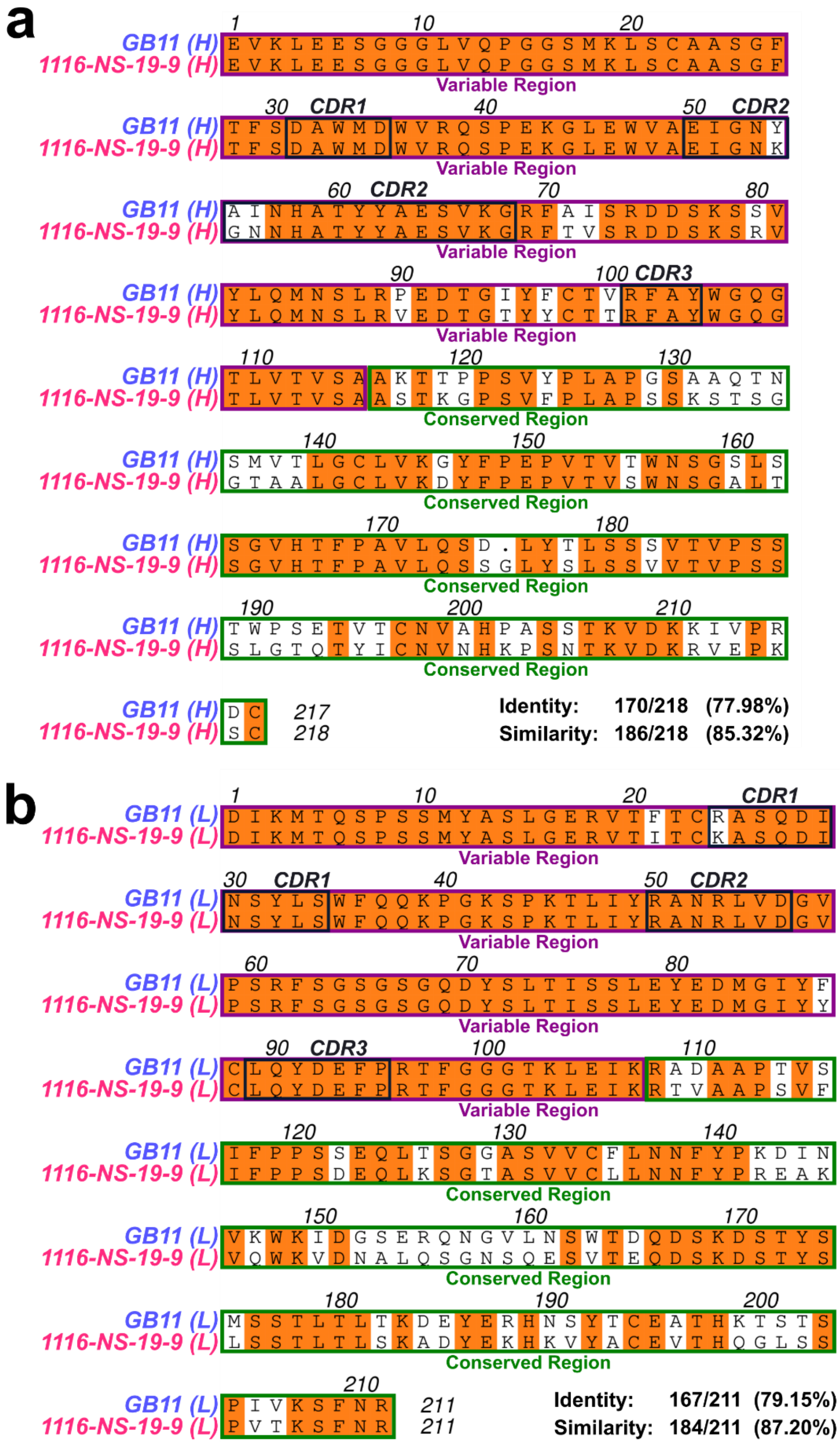


***Figure S3| GB11 shows greater variation in conserved regions than in variable regions compared to 1116-NS-19-9.*** *The sequence alignments of heavy (**a**) and light chains (**b**) of the commercial mAb 1116-NS-19-9 (PDB ID: 6XTG) compared to the novel mAbs GB11 (PDB ID: 9I9H). The alignment is depicting the variable region (purple) as well as the conserved region (green) of the heavy and light chain respectively. The CDR regions are depicted in black. At the end the sequence identity and similarity are displayed.*

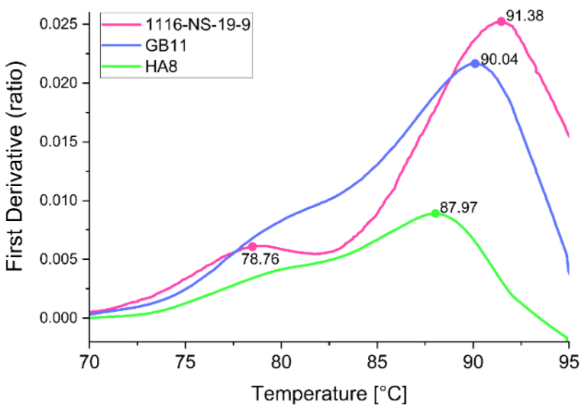


***Figure S4| GB11 and HA8 exhibit similar thermal stability compared to 1116-NS-19-9.*** *Intrinsic fluorescence was recorded at 330 nm and 350 nm while heating the sample from 35 to 95°C at a rate of 3°C/min. Here, the first derivative of the 330/350 nm ratio is shown. The dots depict the melting points T. 1116-NS-19-9 reveals two melting points, called T1 and T2.*

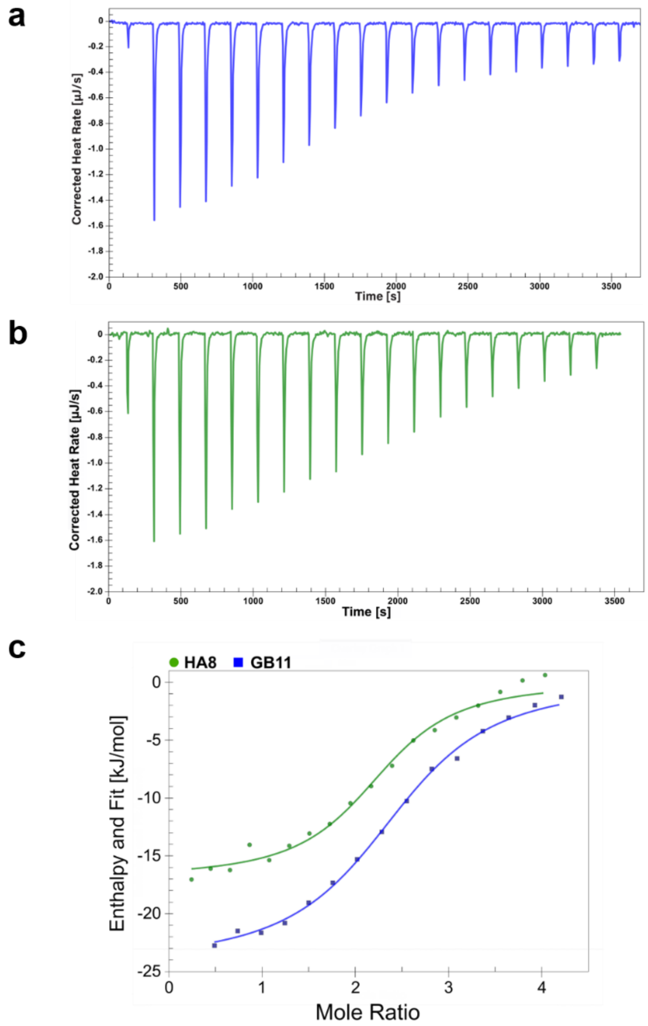


***Figure S5| GB11 and HA8 exhibit similar affinity towards synthetic sLeA.*** *Isothermal titration calorimetry (ITC) of GB11 and HA8 with sLeA. The mAb was in the cell, while sLeA was injected. (**a-b**) Examples for each mAb are shown. GB11 in blue (**a**) and HA8 in green (**b**). Examples of the fitted enthalpy in kJ per mol is presented underneath (**c**).*

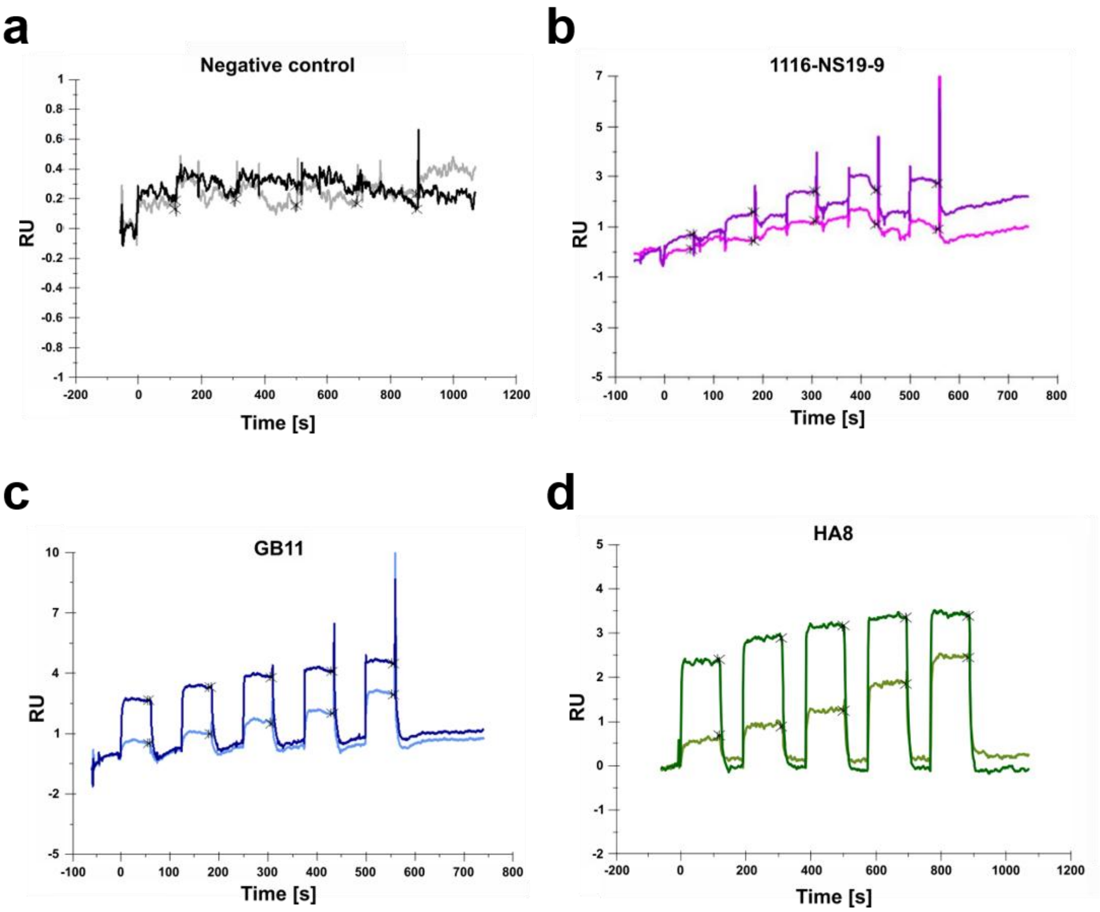


***Figure S6| Raw SPR sensorgrams show concentration-dependent sLeA binding to immobilised mAbs*** *Surface plasmon resonance (SPR) measurements were performed using immobilised mouse monoclonal antibodies (mAbs) and synthetic sLeA as analyte. (**a-d**) Representative raw sensorgrams from one replicate per mAb are shown. Response units (RU) were recorded over time in seconds (s) across increasing concentrations of sLeA. Lower concentrations are shown in lighter shades of colour, higher concentrations in darker shades. The anti-mouse IgG from the immobilisation kit, used as a negative control is shown in grey (**a**). 1116-NS-19-9 is shown in magenta (**b**), GB11 in blue (**c**), and HA8 in green (**d**). RU values were recorded at the indicated time points after washing.*

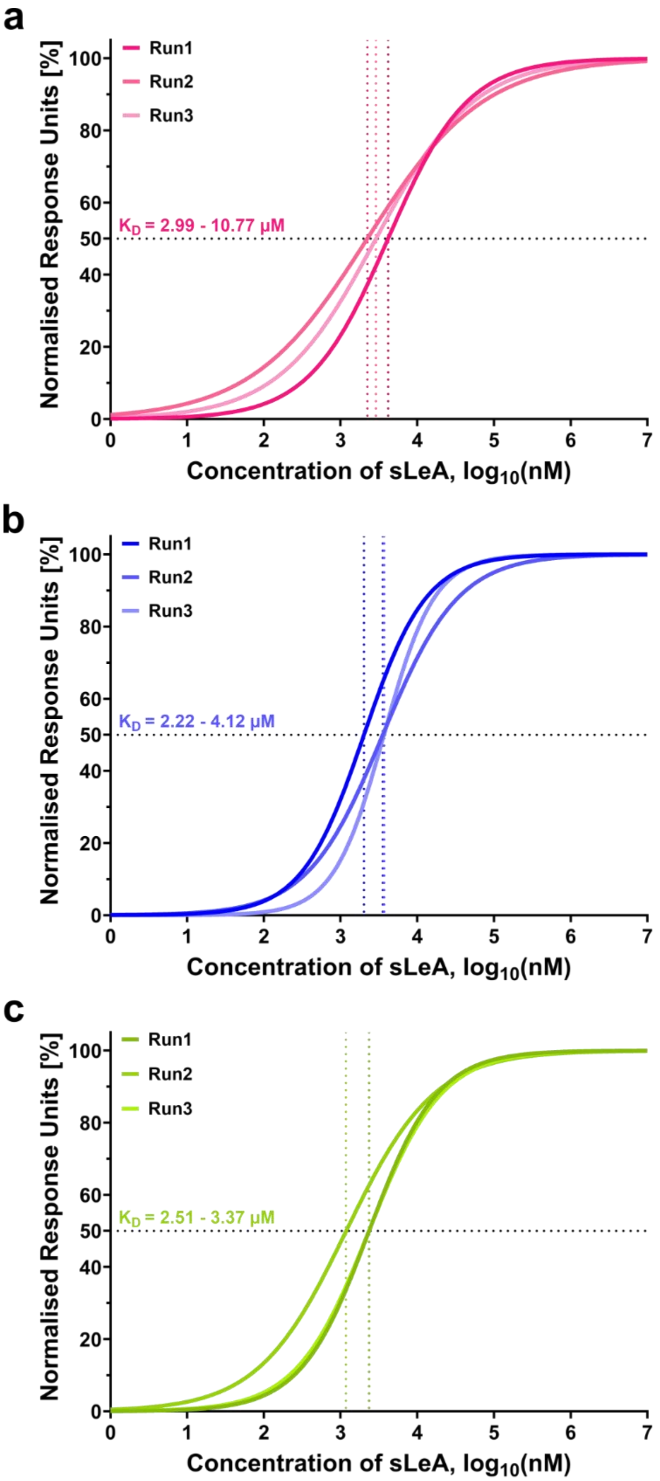


***Figure S7| Fitted SPR data reveals KD ranges in micromolar range for each mAb.*** *SPR measurements were performed using immobilised mouse mAbs and synthetic sLeA as analyte. (**a-c**) Sensorgrams were fitted using a variable-slope, dose–response model in GraphPad Prism (v10.4.2). Three individual runs (run1–3) were performed per mAb and are shown separately. The analysis is presented for 1116-NS-19-9 in magenta (**a**), GB11 in blue (**b**), and HA8 in green (**c**). $K_D$ ranges derived from the Biacore T200 evaluation software 3.2 are indicated in each panel.*

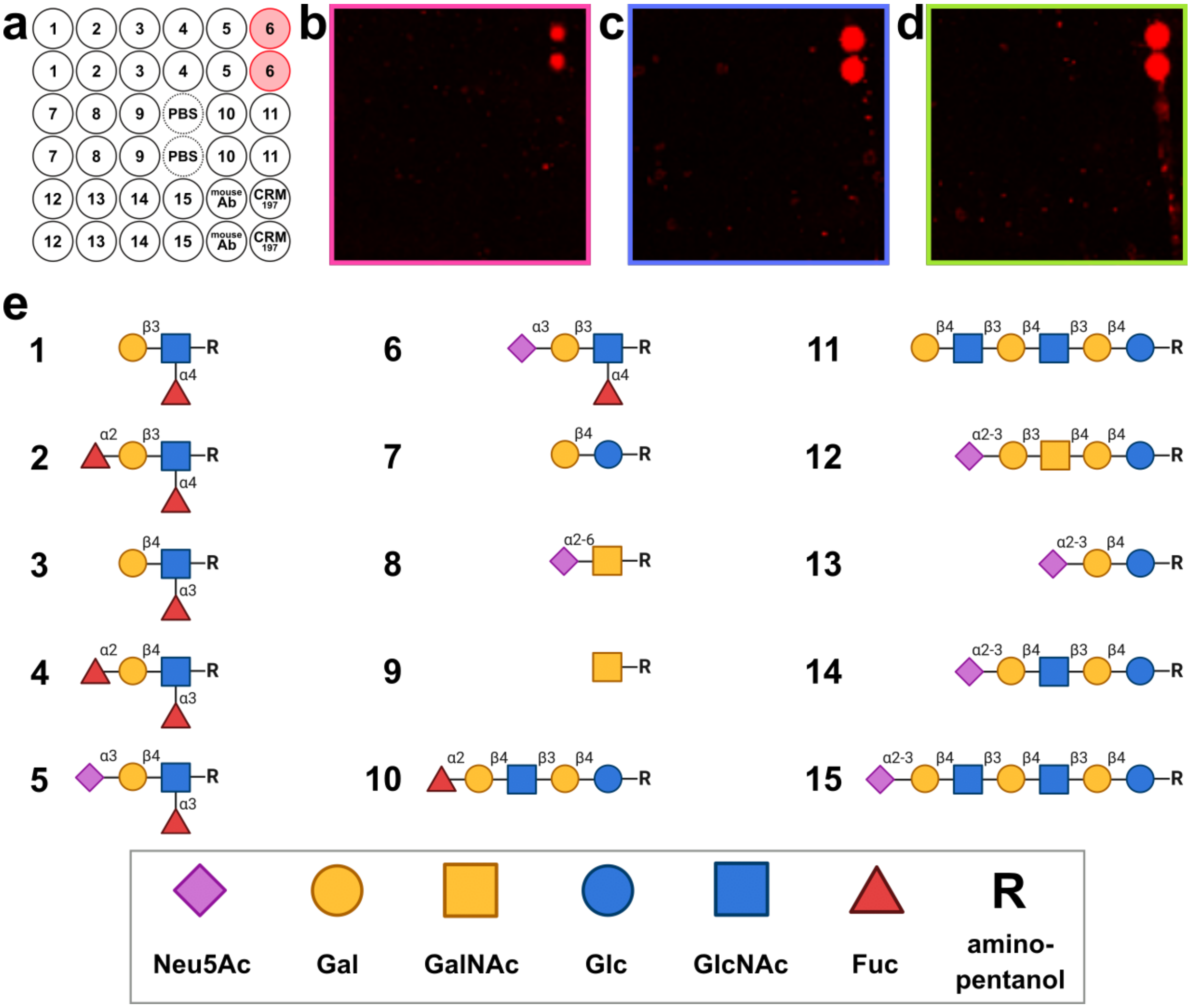


***Figure S8| The mAbs show a high specificity towards sLeA only.*** *(**a**) Printing pattern of the glycan array. The red dots (No. 6) indicate the locations where sLeA is printed. Each number corresponds to the printed glycan structure, which are represented in the "Symbol Nomenclature for Glycans" (SNFG) as shown in (**e**). (**b-d**) Depict one out of four repetitions of the glycan array, demonstrating that the mAbs 1116-NS-19-9 (**b**), GB11 (**c**), and HA8 (**d**) exhibit high antigen specificity, binding exclusively to sLeA. The SNFG representations were created with BioRender.com.*

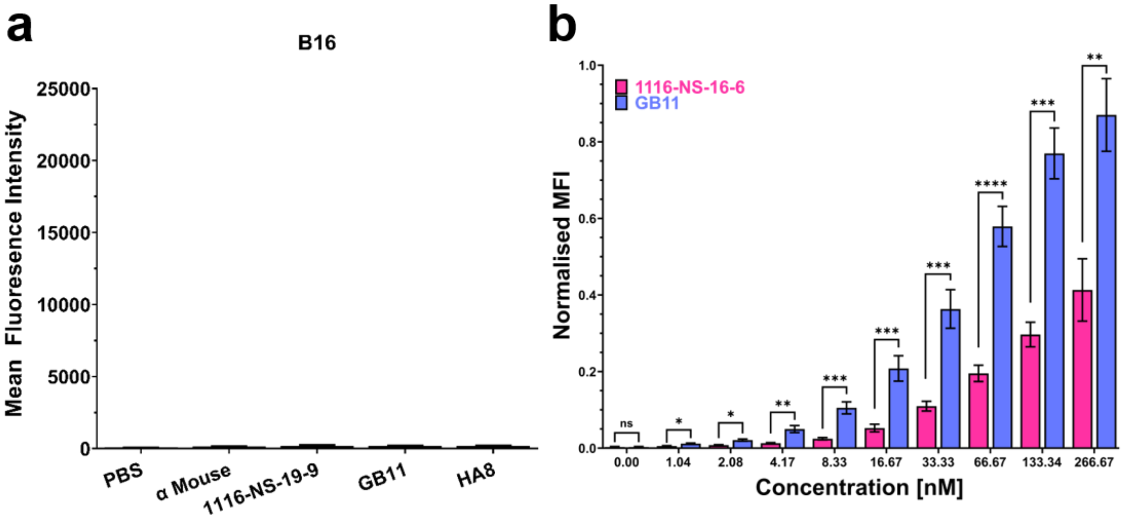


***Figure S9| GB11 binds significantly stronger to sLeA expressing mouse melanoma cells.*** *(**a**) Mean fluorescence intensity (MFI) with standard error of the mean (SEM) for 1116-NS-19-9 (magenta), GB11 (blue), and HA8 (green) on B16 mouse melanoma cells. Five independent assays were conducted for each sample, using 5 µg/mL of the respective mAbs. Statistical analysis was performed using One-Way ANOVA. (**b**) Titration binding assay performed on B16-FUT3+ cells for GB11 (blue) and 1116-NS-19-9 (magenta). The MFI was normalised to the highest measured MFI for each mAb. Error bars represent the SEM. PBS and secondary-only samples are omitted for clarity. Statistical analysis was conducted using One-Way ANOVA (ns = not significant, *p ≤ 0.05, **p ≤ 0.01, ***p ≤ 0.001, ****p ≤ 0.0001).*

***Table S1| X-ray data collection and model refinement statistics.*** *Structures for apo (PDB ID: 9I6Q) and holo (PDB ID: 9I9H) forms were obtained. Values in parentheses refer to the highest resolution shell.*

| | *GB11 apo* | *GB11 sLeA* |
|---|---|---|
| *Wavelength* | 0.9184 | 0.9184 |
| *Reflections* | 469197 | 122337 |
| *Unique* | 40412 | 40947 |
| *Space group* | P4 | R1 |
| *Cell dimensions* | | |
| *a, b, c* | 109.160, 109.160, 40.268 | 39.838, 108.697, 108.920 |
| *α, β, γ* | 90.00, 90.00, 90.00 | 89.60, 87.34, 89.25 |
| *Resolution* | 54.58-1.86 (1.90-1.86) | 26.98-2.90 (3.02-2.80) |
| *Completeness* | 100 (99.6) | 98.2 (93.4) |
| *I/σ(I)* | 7.7 (0.9) | 2.6 (0.6) |
| *CC1/2* | 0.996 (0.376) | 0.915 (0.189) |
| *Rmerge* | 0.202 (2.871) | 0.327 (2.035) |
| *Multiplicity* | 11.6 (11.6) | 3 (3.2) |
| ***Refinement*** | | |
| *Resolution* | 48.82-1.68 | 26.70-2.96 |
| *Rwork/Rfree* | 0.205/0.240 | 0.210/0.248 |
| *Number of monomers* | 1 | 4 |
| *Average B factors* | 23.74 | 62.86 |
| *GB11* | 22.879 | 63.1 |
| *Ligand* | - | 78.83 |
| *Solvent* | 37.523 | 52.2 |
| *RMS Z-score* | | |
| *Bonds* | 0.758 | 0.441 |
| *Angles* | 0.973 | 0.759 |
| *Ramachandran* | | |
| *Favoured* | 413 (97.18%) | 1588 (93.41%) |
| *Allowed* | 11 (2.59%) | 100 (5.87%) |
| *Disfavoured* | 1 (0.24%) | 15 (0.88%) |

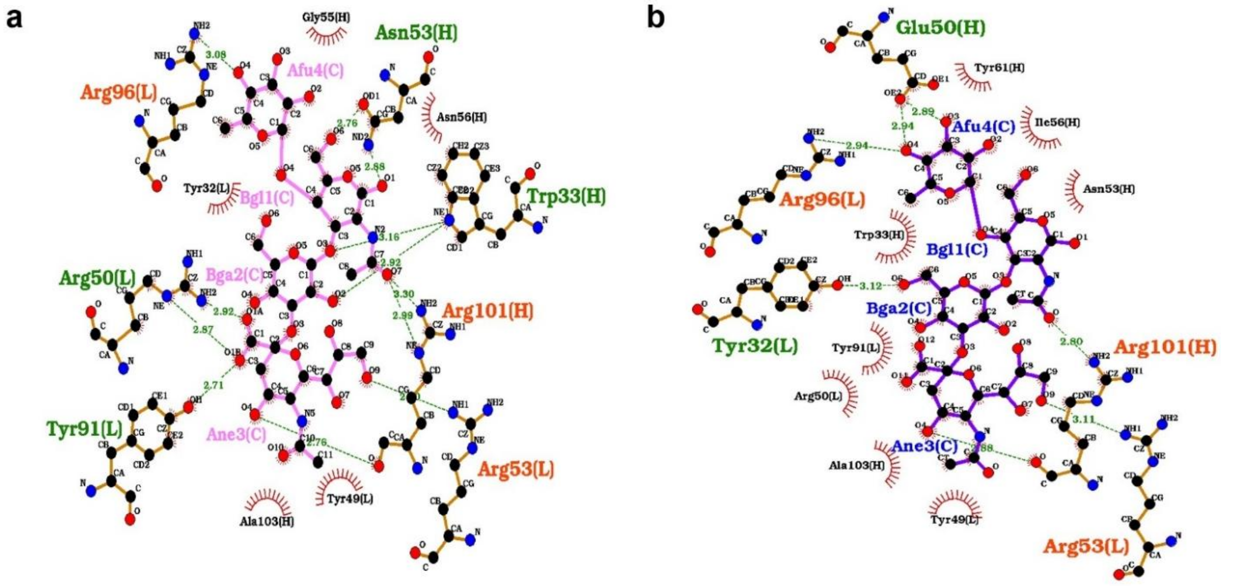


***Figure S10| Ligplot results of the minimised docked complex of 1116-NS-19-9 and GB11 from AutoDock Vina.*** *(**a**) The plot represents the h-bond distance and interacting residues of the complex 1116-NS-19-9 with sLeA and in (**b**) for the complex of GB11 with sLeA. The binding sites residues forming h-bond interaction in both the complexes are named in orange-red colour and residues that are specific for each of the mAbs are named in olive green.*

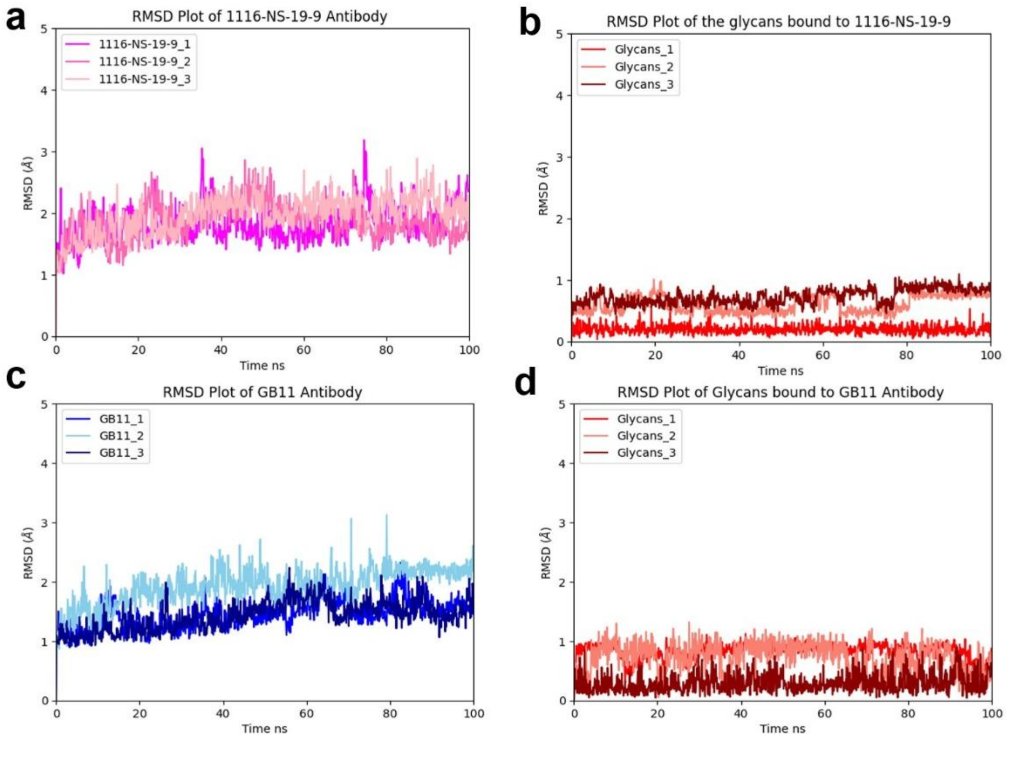


| 1116-NS-19-9_Complex | Average RMSD |
|---|---|
| 1116-NS-19-9_replicate_1 | 1.891 ± 0.324 |
| 1116-NS-19-9_replicate_2 | 1.703 ± 0.485 |
| 1116-NS-19-9_replicate_3 | 1.890 ± 0.319 |
| Glycans_1 | 0.197 ± 0.306 |
| Glycans_2 | 0.598 ± 0.374 |
| Glycans_3 | 0.725 ± 0.291 |

| GB11_Complex | Average RMSD |
|---|---|
| GB11_replicate_1 | 1.447 ± 0.266 |
| GB11_replicate_2 | 1.920 ± 0.272 |
| GB11_replicate_3 | 1.425 ± 0.287 |
| Glycans_replicate_1 | 0.830 ± 0.307 |
| Glycans_replicate_2 | 0.726 ± 0.235 |
| Glycans_replicate_3 | 0.295 ± 0.256 |

***Figure S11| SLeA binding stabilises GB11 and 1116-NS-19-9 complexes.*** *(**a**) The last 100 ns averaged backbone atom RMSD values of all three replicates of 1116-NS19-9 exhibit significant overlap, indicating consistency across the replicates. (**b**) The averaged backbone atom RMSD of the tetrasaccharide bound to 1116-NS19-9 remained consistently lower than that of 1116-NS19-9, suggesting a stable retention of sLeA within the binding pocket. (**c**) The last 100 ns averaged backbone atom RMSD values of all three replicates of GB11 show good overlap within the admissible range of 3.0 Å.(**d**) sLeA bound to GB11 exhibited a backbone atom deviation with an RMSD value of less than 1.0 Å across all three replicates, indicating stable ligand binding.*

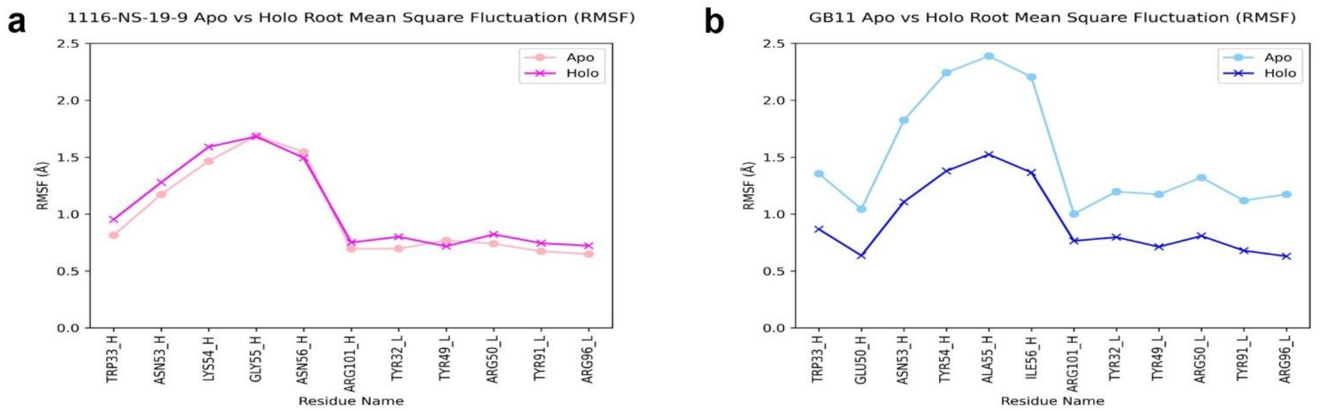


***Figure S12| Tetrasaccharide binding reduces binding site flexibility in GB11 but not in 1116-NS-19-9.*** *The RMSF values were computed and averaged across replicates, focusing on binding site residues of 1116-NS-19-9 and GB11. (**a**) 1116-NS19-9 exhibited minimal fluctuations in the RMSF values of the binding site residues in both the apo and holo forms. (**b**) In contrast, GB11 when comparing its apo and holo forms, showed reduced fluctuations in the binding site residues in the holo form upon tetrasaccharide binding, suggesting increased backbone rigidity.*

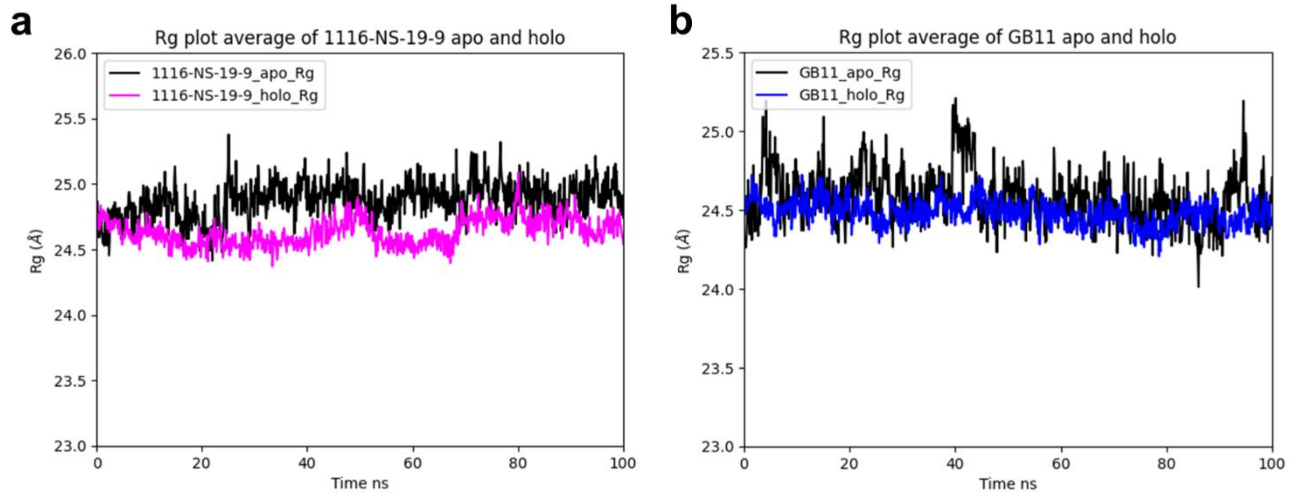


***Figure S7| The average radius of gyration (Rg) apo and holo mAb.*** *The plot reveals that the protein structure maintains a high degree of compactness, with a Rg value consistently fluctuating within the acceptable range of < 1 Å for both 1116-NS-19-9 (**a**) and GB11 (**b**) in both their apo and holo forms. However, in the GB11 holo form the compactness looks well preserved.*

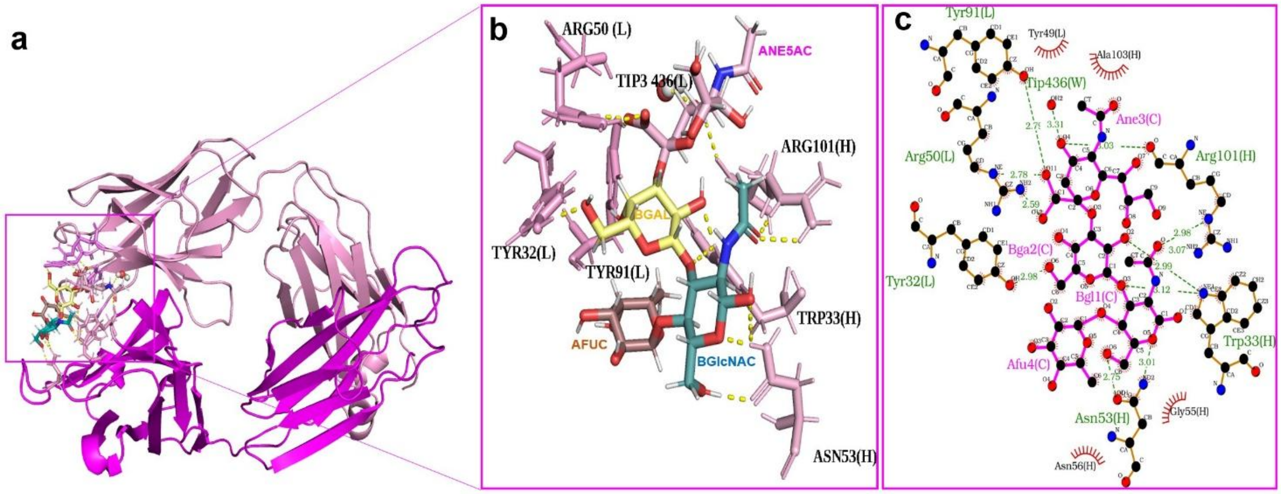


***Figure S84| 1116-NS-19-9 binding site for sLeA with lowest interaction energy score.*** *(**a**) Binding site of the 1116-NS-19-9 mAb. (**b**) Displaying the residues engaged in forming polar H-bond contacts with the glycan residues (**c**) Its 2D Ligplot showing the H-bond distance between the glycans and the key residues.*

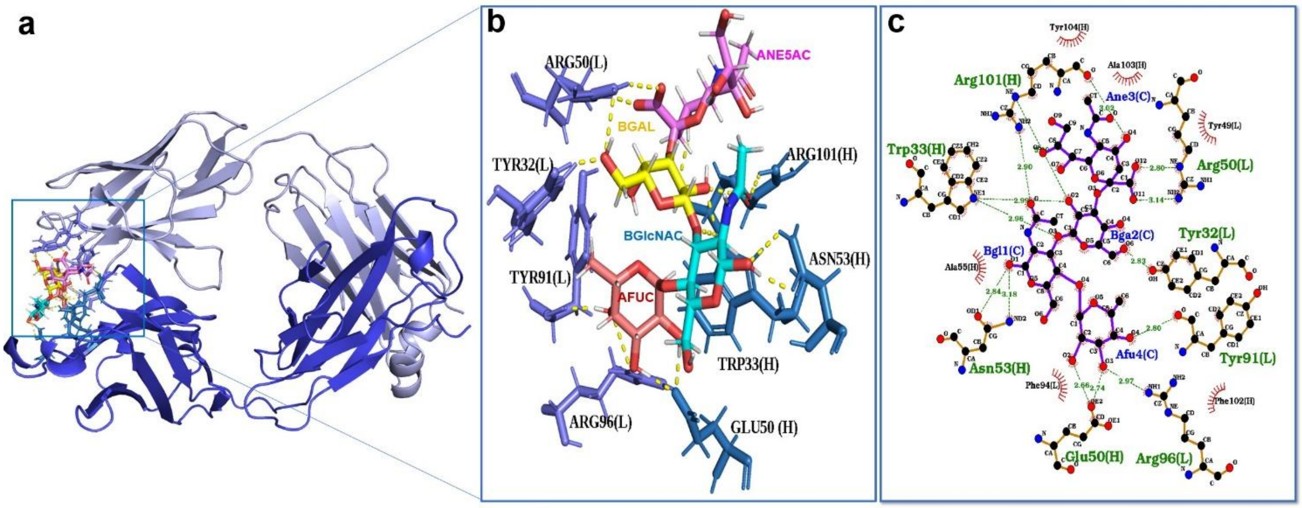


***Figure S15| GB11 binding site for sLeA with lowest interaction energy score.*** *(**a**) Binding site of the GB11 mAb. (**b**) It reveals the vital residues involved in forming polar H-bond contacts with the glycan residues. (**c**) A 2D Ligplot provides a visualization of the H-bond distances between the glycans and the critical residues.*

***Table S2| Chemical Shift Assignment of sialyl Lewis A.*** *The table contains the chemical shift values in ppm for $^{1}H$ and $^{13}C$ atoms of sLeA. The commercially available sLeA comprises a mixture of α- and β-GlcNAc anomers, attributable to the presence of a free OH-group at the CH1 of GlcNAc. Consequently, resonances corresponding to both α- and β-GlcNAc are observed, and are represented with α- and β- respectively.*

| | Position | $^{1}H$ [ppm] | $^{13}C$ [ppm] |
|---|---|---|---|
| α-GlcNAc | 1 | 5.0450 | 90.9749 |
| | 2 | 4.0695 | 53.9827 |
| | 3 | 4.1043 | 74.1583 |
| | 4 | 3.6809 | 72.3631 |
| | 5 | 3.9112 | 71.4154 |
| | 6 | 3.8045 | 59.6534 |
| | NAc-Methyl | 1.9724 | 22.1618 |
| β-GlcNAc | 1 | 4.6503 | 94.7612 |
| | 2 | 3.7908 | 56.7851 |
| | 3 | 3.9882 | 76.1052 |
| | 4 | 3.6646 | 72.2969 |
| | 5 | 3.4914 | 75.5419 |
| | 6 | 3.8805 | 59.7005 |
| | NAc-Methyl | 1.9724 | 22.1618 |
| Gal-α-GlcNAc | 1 | 4.4921 | 102.7904 |
| Gal-β-GlcNAc | 1 | 4.4774 | 102.7944 |
| Gal | 2 | 3.4331 | 68.7804 |
| | 3 | 3.9811 | 75.6270 |
| | 4 | 3.8384 | 66.8756 |
| | 5 | 3.4594 | 74.6491 |
| | 6 | 3.6229 | 61.4691 |
| Fuc | 1 | 4.9419 | 97.9594 |
| | 2 | 3.7234 | 67.8117 |
| | 3 | 3.8126 | 69.1054 |
| | 4 | 3.7106 | 71.9605 |
| | 5 | 4.8123 | 66.8383 |
| | 6 | 1.1033 | 15.3380 |
| Neu5Ac | 3-1 | 2.7012 | 40.0003 |
| | 3-2 | 1.6935 | 40.0008 |
| | 4 | 3.6062 | 68.4542 |
| | 5 | 3.7792 | 51.6653 |
| | 6 | 3.5486 | 72.6980 |
| | 7 | 3.5445 | 67.9888 |
| | 8 | 3.7808 | 71.8305 |
| | 9-1 | 3.7492 | 62.2379 |
| | 9-2 | 3.5796 | 62.2374 |
| | NAc-Methyl | 1.9596 | 22.0634 |

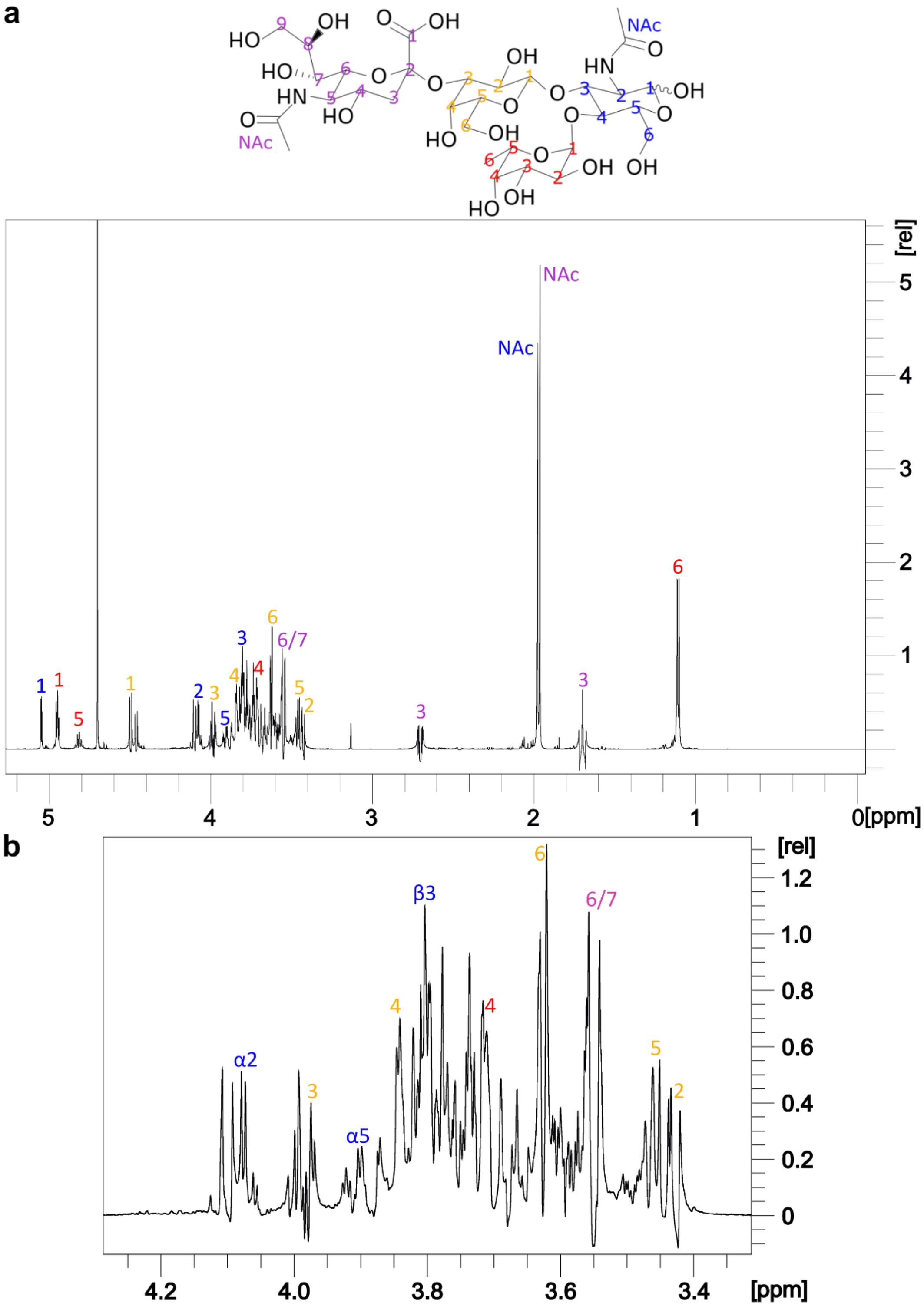


***Figure S16| [1]H NMR spectrum and assignment of sLeA.*** *(**a**) On the top is the structure of sLeA, for the assignment each carbon atom is represented with numbers in the colours of the monosaccharides in the SNFG representation, GlcNAc (blue), Fuc (red), Gal (yellow) and Neu5Ac (purple). These numbers are used on the respective resonances of sLeA in the [1]H-spectrum. (**b**) The zoomed-in [1]H spectrum spans the range between 4.2 and 3.4 ppm. Anomers of α- and β-GlcNAc are represented with α and β respectively. The assignment presented herein is relevant to the STD NMR analysis.*

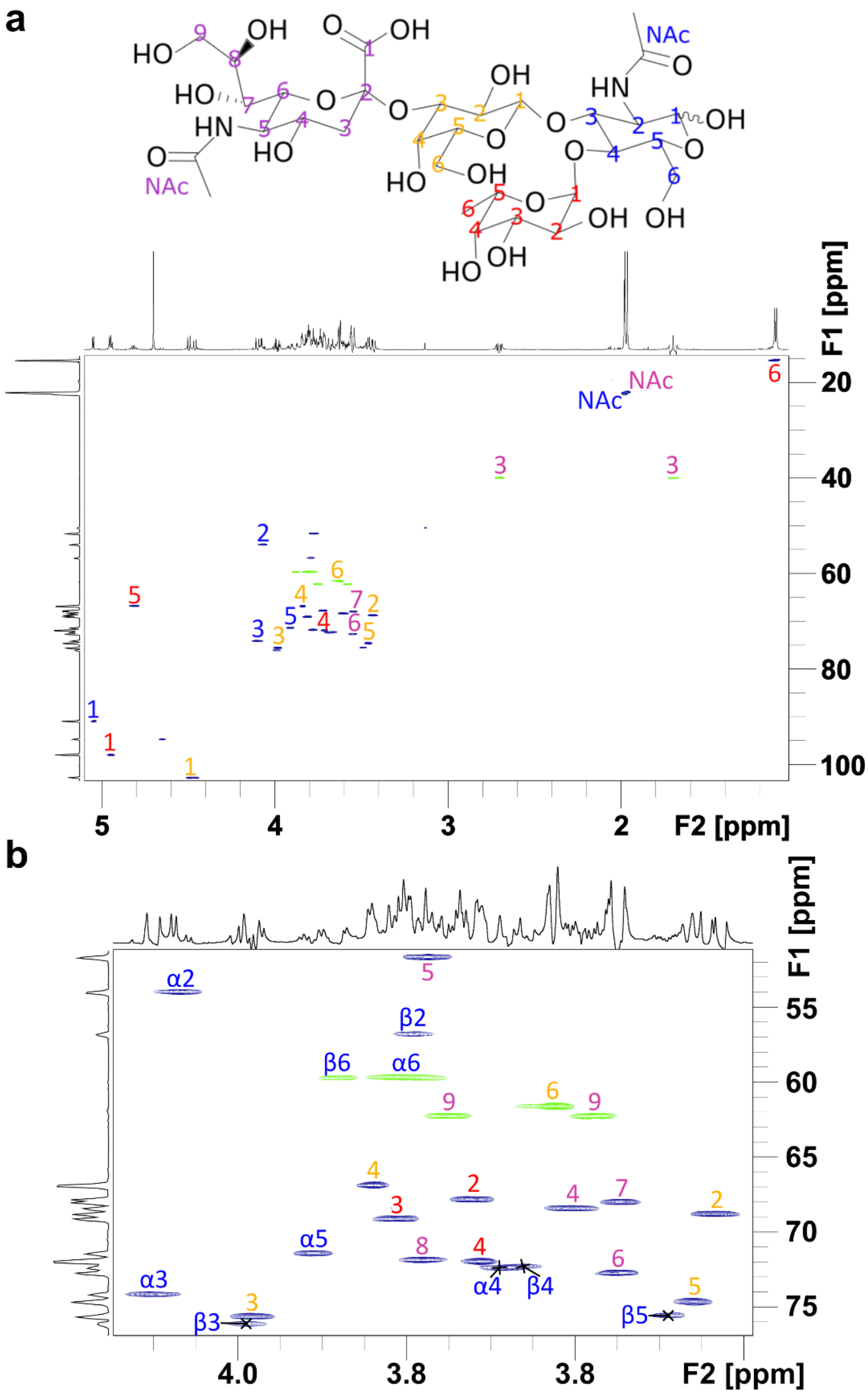


***Figure S97| $^{1}$H-$^{13}$C HSQC and assignment of sLeA.*** *(**a**) On the top is the structure of sLeA. For the assignment each carbon atom is represented with numbers in the colours of the monosaccharides in the SNFG representation, GlcNAc (blue), Fuc (red), Gal (yellow) and Neu5Ac (purple). These numbers are used on the respective resonances of sLeA in the $^{1}$H-$^{13}$C HSQC. (**b**) $^{1}$H-$^{13}$C HSQC zoomed in between 55 and 75 ppm for F1 ($^{13}$C) and 4.2 and 3.6 ppm for F2 ($^{1}$H). Anomers of α- and β-GlcNAc are represented with α and β respectively. In (**a**) we present the assignment used for the STD NMR analysis and in (**b**) respective assignment for α- and β-GlcNAc.*

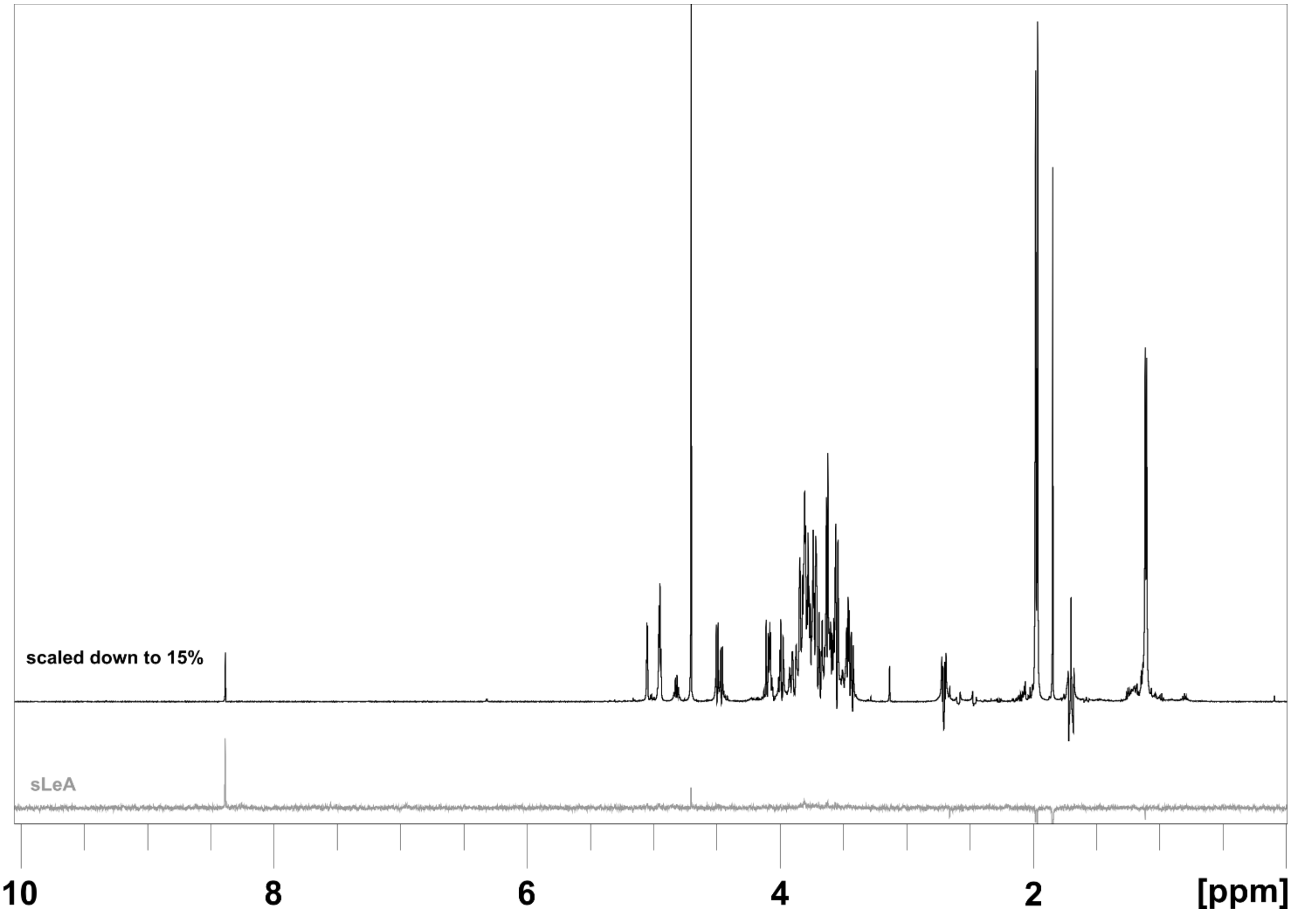


***Figure S108| In the absence of a protein, the STD spectrum of sLeA exhibits minimal saturation.*** *Protein saturation was achieved using low-power Gaussian-shaped pulses at 8.25 ppm, with a total duration of 2 seconds. The grey spectrum illustrates the STD spectrum of sLeA in the absence of a protein.*

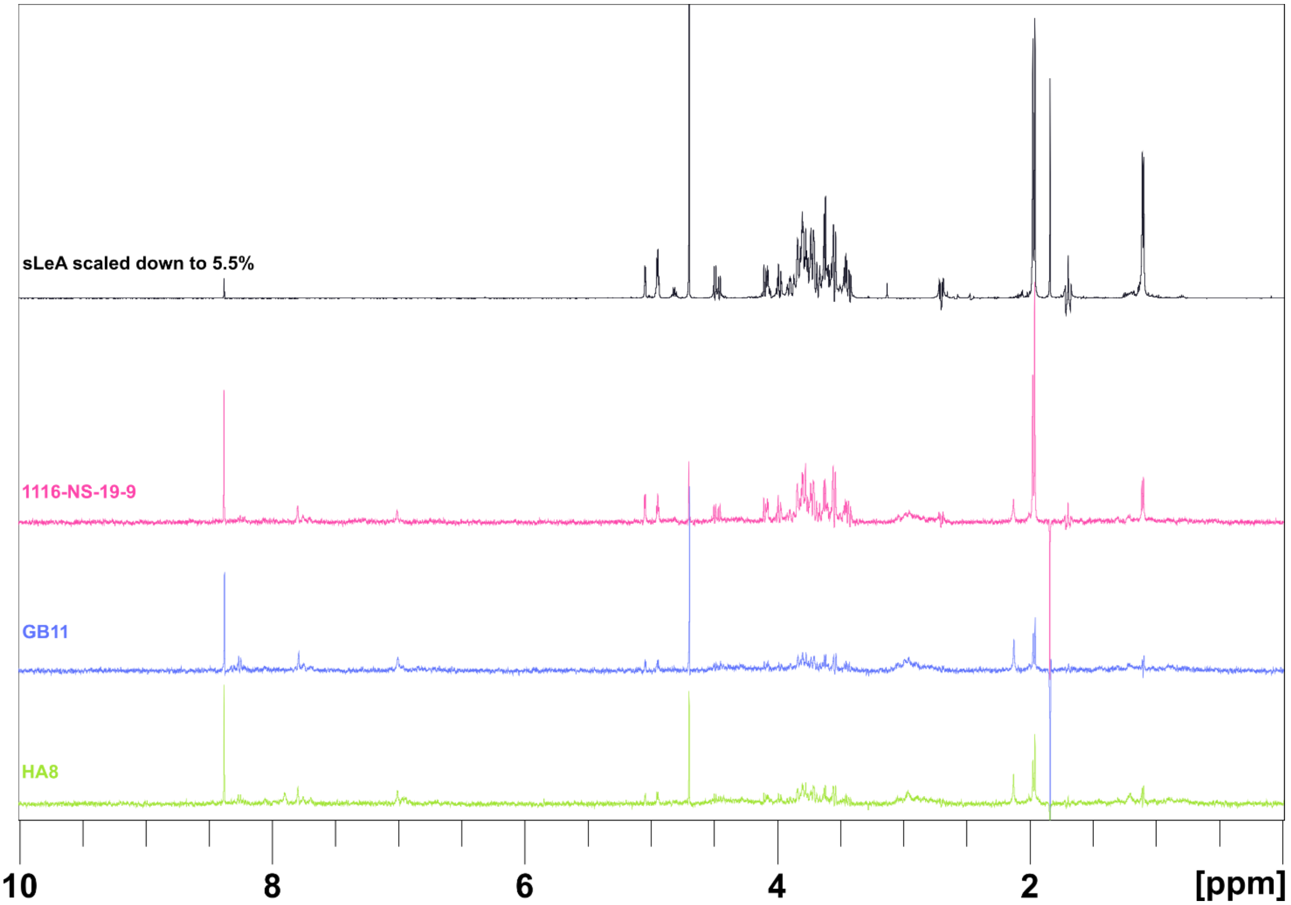


***Figure S119| The STD spectra of sLeA demonstrate interaction with all three mAbs.*** *At the top, the $^{1}$H reference spectrum of 200 µM sLeA, reduced to 5.5%, is depicted in black. In the lower section of the image, the STD spectra of 200 µM sLeA are presented, recorded concurrently with 1116-NS-19-9 (magenta), GB11 (blue), or HA8 (green) at a protein concentration of 6.7 µM each. It is evident that all three monoclonal antibodies bind to sLeA. Furthermore, 1116-NS-19-9 exhibits on average higher STD effects, suggesting differences in binding kinetics compared to GB11 and HA8.*

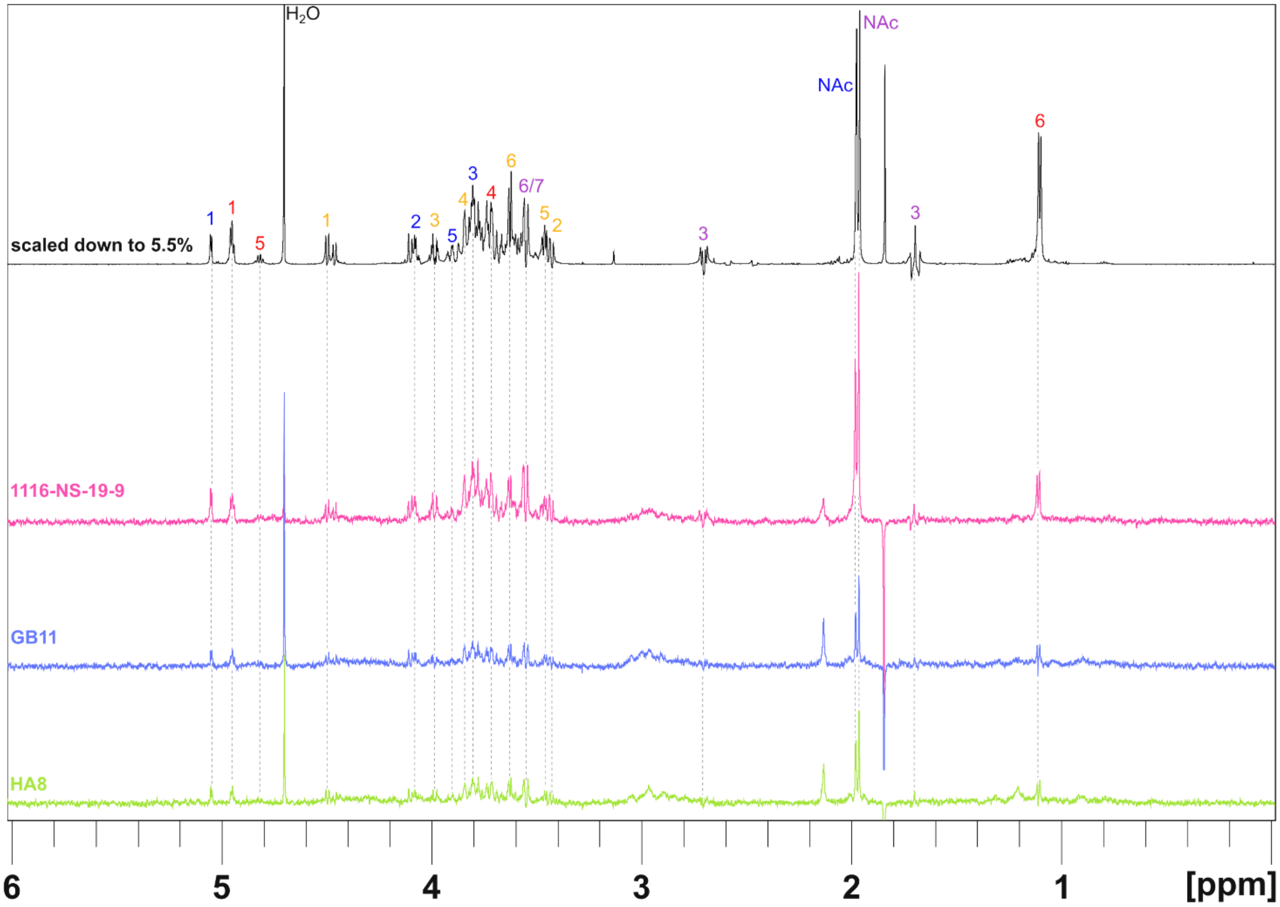


***Figure S20| The STD spectra depict the resonances involved in the interaction between sLeA and the respective antibody**. The $^{1}H$ reference spectrum of 200 µM sLeA, reduced to 5.5%, is illustrated in black at the top. Each hydrogen atom utilized for STD effect measurement is labelled with numbers corresponding to the monosaccharide colours in the SNFG representation: GlcNAc (blue), Fuc (red), Gal (yellow), and Neu5Ac (purple). In the lower section of the image, the STD spectra of 200 µM sLeA are displayed, recorded in conjunction with 1116-NS-19-9 (magenta), GB11 (blue), and HA8 (green) at a protein concentration of 6.7 µM each.*

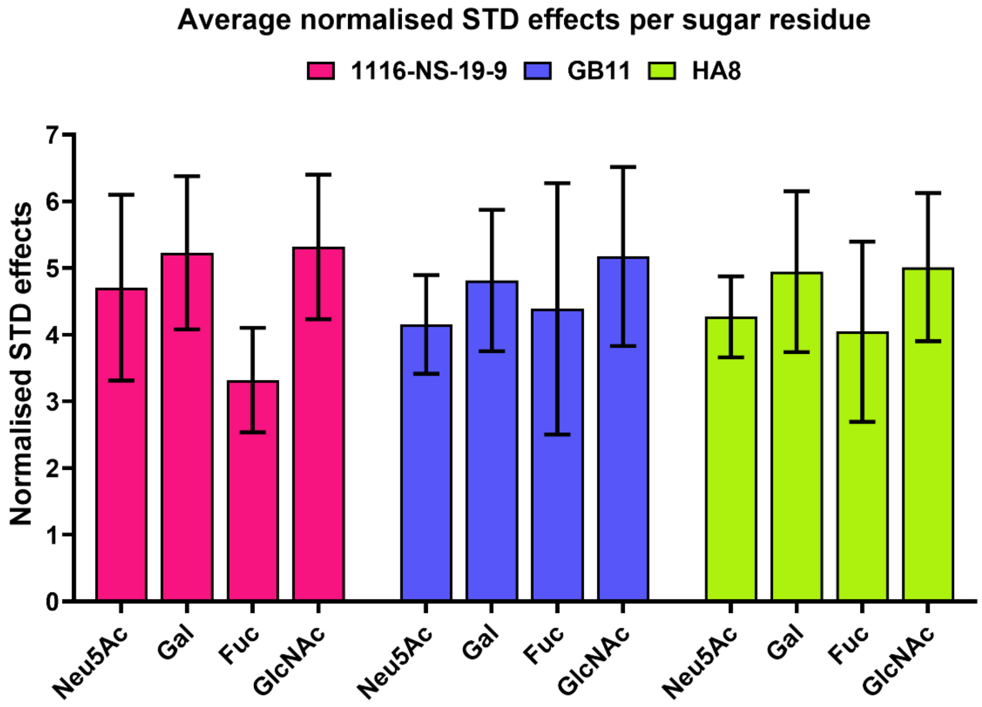


***Figure S2112| GB11 and HA8 show more uniform STD NMR saturation across sLeA glycan units than 1116-NS-19-9.*** *Average normalised STD effects per sugar residue are shown for each antibody: 1116-NS-19-9 in magenta, GB11 in blue, and HA8 in green. GB11 and HA8 display more uniform saturation across all four glycan units of sLeA, whereas 1116-NS-19-9 shows notably weaker saturation at the fucose residue, indicating less engagement of this part of the epitope.*
